# Sir John Murray’s H.M.S. Challenger Sedimentary Deposits Collection at the Natural History Museum, London

**DOI:** 10.1101/2025.03.21.644325

**Authors:** C. Giles Miller, Monique M. H. Jouet-Sarkany

**Affiliations:** Natural History Museum, Cromwell Road, London SW7 5BD

**Keywords:** Sir John Murray, H.M.S. Challenger, marine sediment, Natural History Museum, Edinburgh

## Abstract

The Natural History Museum holds 4,718 bottles, tubes, boxes, and slides containing sediment and derived preparations from 80% of the 504 listed marine collecting localities of H.M.S. Challenger (1873-1876). The sediment collections include manganese nodules, pumice, and various marine vertebrates and invertebrates, predominantly the Foraminifera from all the major oceans of the world. A complete online database of the collection is presented for the first time to include all of the associated collecting and preparation data from the oceanic dredges, trawls, soundings and anchor mud collections alongside a set of previously undocumented shallower water collections that were not included in the Challenger Deep Sea Deposits Report of Murray and Renard (1891). The dataset allows assessment of various other Challenger sediment derived collections dispersed worldwide in museums and private collections. A historical account of the collection follows; collecting and studying onboard, studying in Edinburgh post cruise, donation by Sir John Murray’s family to the British Museum (Natural History) and accounts of the collection’s historical housing at the museum. The historical context and presented dataset are intended to aid future users of the collection to select suitable materials for study or display. A review of previous scientific and exhibition uses of the collection is followed by suggestions for the future in terms of storage and potential for illuminating the debate on anthropologically driven climate, biodiversity and atmospheric changes.

## Introduction

Much has been written and continues to be written about the famous H.M.S. Challenger Expedition (1872-1876) that circumnavigated the globe with the aim of studying the biology, chemistry and physics of the ocean and its bottom sediments (eg Linklater, 1972; Rice, 1972, 2001; McDougall, 2019; Jones, 2023 and Wood, 2025). Published and unpublished diaries written by some of the crew members and scientific staff (e.g. Wild, 1878 and Rehbock, 1993, diary of Assistant Ship’s Steward Joseph Matkin) also provide a historical narrative that puts this groundbreaking cruise into perspective. The 50 bound volumes representing various aspects of the cruise remain key to unlocking the scientific and historical significance of collections from the voyage. McDougall (2019, Chapter 2) gives a good brief biography of the members of the scientific party including the scientific leader of the expedition Charles Wyville Thompson, Chemist John Buchanan, Artist John James Wild and Naturalists Rudolf Von Willemoes-Suhm, Henry Moseley and John Murray. Born in 1841 in Ontario, Canada, Murray moved back to Scotland where his Scottish grandfather appointed him curator of his Natural History Museum while still at school. He was later encouraged to study medicine at the University of Edinburgh but never finished his degree. Instead, he got a job as surgeon on whaling ship where he made observations of natural history and studied bottom sediments which no doubt helped when he was recommended to Wyville Thompson for a place on the Challenger Expedition. He was responsible for studying the sediments and other geological particles dredged from the ocean depths. After the cruise he worked with Wyville Thompson in the Challenger office in Edinburgh distributing the collections and encouraging authors to finish their contributions. When Wyville Thompson stepped back from his duties due to ill health, he oversaw publication of the 50 Challenger volumes, contributing to three of them but most significantly the penultimate volume on Deep Sea Deposits (Murray and Renard, 1891) that also included reference to material collected by subsequent investigations by the India Rubber Company, Telegraph Works Company and Norwegian, Italian, French, German, and American Expeditions.

This paper does not include details of these subsequent collections but focuses on sedimentary material collected by H.M.S. Challenger between 1873 and 1876 that is housed at the Natural History Museum and referenced in the Deep Sea Deposits volume (Murray and Renard, 1891). Originally the volume set out to describe material collected from depths greater than 100 fathoms which excludes some of the shallower water samples taken near to harbouring spots such as Kerguelen Island, Hong Kong and Sydney. Sometimes these were not given official station collection numbers but they are also included in this dataset as are other collections made between the officially numbered Challenger collecting stations. Associated with the collection are several hundred plankton slides that were collected from surface plankton tows but referenced in Murray and Renard (1891) and Murray (1876) but these are not included in the dataset as they will be the subject of a future paper on the plankton collections. It should also be noted that the focus of this paper is the sediments in John Murray’s collection rather than macro and micro-organisms collected at the same time and made the subject of most of the 50 published volumes. Likewise, any sediment subset collections derived from Sir John Murray’s Sediment Collection but now housed away from the Murray Collection at the Natural History Museum (for dispersal details see Lingwood 1981) are also out of scope for this study.

The Natural History Museum, London (NHM) was originally housed with the collections of Hans Sloane in Bloomsbury where he founded the British Museum in 1756. The Natural History Collections were moved to a new site in South Kensington in 1881 where the British Museum (Natural History) (BM(NH)) was founded. The museum became independent of the British Museum in 1992 and was renamed the Natural History Museum (NHM). In this paper, date context will be used to reference the historical naming of the institution which has housed the collection in several locations both onsite in South Kensington and offsite where it is currently housed.

This contribution accompanies the first detailed dataset of Sir John Murray’s Challenger Sediment Collection outlining 4,718 individual items with verbatim label transcription and details of how they were collected and prepared for study. This paper also covers a brief history of storage location conditions, a review on the history of use as well as suggestions for future use. The listings are intended to help other museums and collectors possessing widely dispersed Challenger materials, assess and provide some context to their Deep Sea Deposit and other sediment holdings. It also serves to encourage the use of Challenger sediment collections at The Natural History Museum and elsewhere by outlining the historical aspects of how the collections were made and used.

## Materials and Methods

### Collecting on board H.M.S. Challenger

#### Preparing to collect

To collect from the depth of the oceans, 25,000 fathoms (45.72km) of boltrope were provided for the dredge, as well as 10,000 fathoms of 3-inch rope, 10,000 fathoms of 2.5-inch rope and 5,000 fathoms of 2-inch rope (Wyville Thomson, 1877). By all accounts, the deck where they were kept was very crowded (Matkin in Rehboch, 1993). 200 cases making up a total of 2,300 jars which were all 9 inches high but with different opening diameters were taken on board prior to departure. These were manufactured by E Breffitt and Co of Upper Thames Street, London (Wyville Thomson, 1877). These were known as ‘drop bottles’ (Wyville Thomson, 1877), apparently manufactured for holding sweetmeats of various kinds (Figure 1a) but the listing of glassware taken on board in 1873 (Tizard et al. 1885, p.43) refers to them as ‘rock bottles’, a term we have used in this paper and in the associated dataset. Aside from these bottles which were particularly well suited for large quantities of sediments or animals, thousands of smaller stoppered and corked test tubes of different sizes were also packed away (Wyville Thomson, 1877).

**Figure 1.**
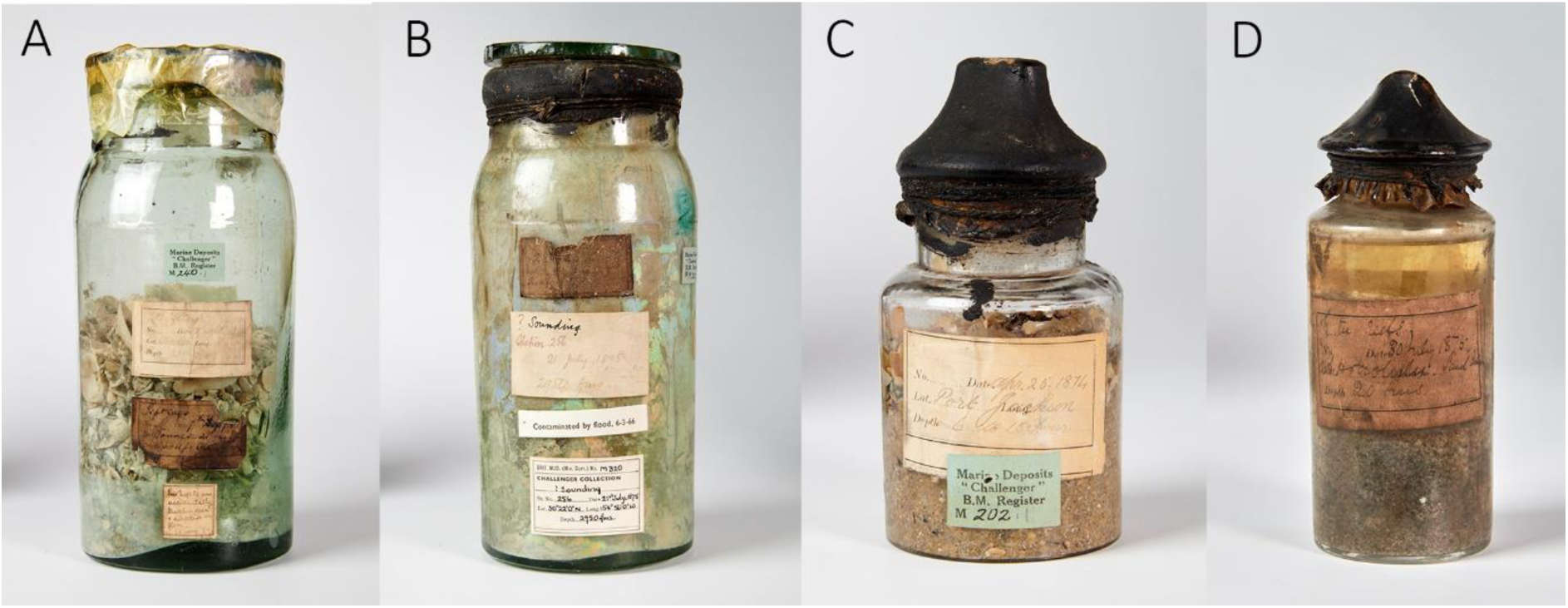
A selection of specimen bottles. A. Rock bottle with sediment and visible shell material NHMUK M.240(1). B. Rock bottle showing iridescence due to glass delamination NHMUK M.320(1). C. Bottle still sealed NHMUK M.202(1). This is the most common style of bottle in the collection, but most are unsealed. D. Bottle with preserved liquid probably spirit NHMUK M.323(2).

Many of these jars and tubes were used to store the deep-sea deposits that were brought up to the surface and a number of them have remained unopened since the day they were sealed, over 150 years ago (Figure 1b). A very small number still have remains of liquid, originally alcohol, that was put in every bottle to preserve the collected specimens and sediments (Figure 1c). The legendary Victorian fastidious and industrious approach to collecting was de rigueur on HMS Challenger as dictated by the Scientific Director of the Expedition, Wyville Thomson (Wyville Thomson, 1877). Every item was carefully recorded in a regular journal of the day to be kept by the Scientific Director. In addition, each member of the Scientific Staff was provided with a notebook to enter their observations and proceedings, also to be passed on to the Scientific Director. ‘Meticulous’ was the operative word on this expedition, to produce results that would be true and useful.

The Preface to Murray and Renard (1891, p. xi) states that “During the Voyage of the Challenger, the collection, examination, and preservation of all the samples of deposits were undertaken by Mr. Murray.” Several publications give precise details of how these samples were collected (eg Wyville Thomson, 1877; Tizard et al. 1885, Murray et al. 1885) so the intention here is not to replicate that detail but to summarise methods and suggest where variations in either the collecting or the initial study may influence the subsequent use of the collection. In general, there are seven methods of collection represented.

#### Sounding

The Narrative of the Cruise (Tizard et al. (1885), p. 59-66) gives a detailed account of the apparatus used and method for sounding. Both the Hydra and the Baillie sounding machines (Fig. 2a, b) were used over the voyage with the latter preferred towards the end. The general principle for sounding was that a weighted tube was lowered to the sediment surface. The speed at which it dropped was carefully monitored and the total length of rope recorded at the point that the rate slowed dramatically at the presumed point that the tube hit the bottom and the weight drove it part way into the sediment. The narrow tube that continued into the sediment on impact had a butterfly valve that snapped closed as the rope was hauled to the surface, preventing sediment loss. The tube containing the top layer of the sediment was then hoisted to the surface, leaving the weight on the bottom and any sediment trapped in the tube recovered on deck. Depending on the nature of the sediment and the probability that the sounding tube would penetrate easily, sometimes a tube was employed without the butterfly valve.

**Figure 2.**
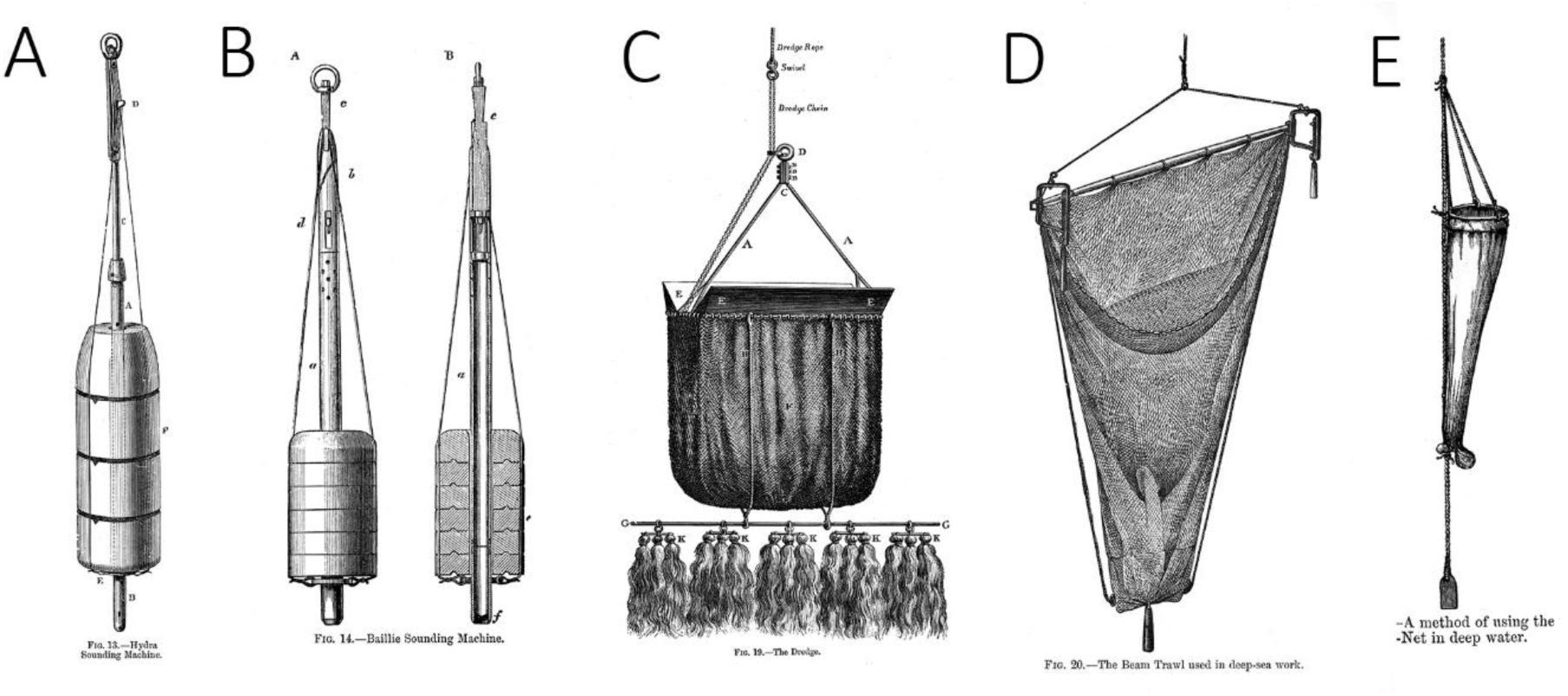
Sample collecting apparatus (from Tizard et al. 1885). A. Hydra Sounding Machine. B. Baillie Sounding Machine. C. Dredge. D Trawl. E. Plankton net.

The size of the sounding samples in the collection indicate that often only a small amount of sediment from soundings was recovered and collected. In the first 6 months of the voyage until July 1873, the tube was very narrow and only a small amount of sediment was able to be collected but this was replaced with a 2 inch diameter tube (Murray, 1876). On occasions the labels with the collection indicate that the material is from either the top or the bottom of the tube (eg M.312). However, it should be assumed that most of the time, the sediment collected represents a mixture of sediments over a shallow depth. Murray (1876, p. 472) reported that sometimes the penetration was as much as 18 inches but also that, “When the tube arrives on board the contents are carefully removed, and the colour, extent, and arrangement of the section is noted. A portion is washed several times in distilled water and dried, a portion is submitted to a rough analysis, and the remainder, if any, is preserved in spirit.” The deepest soundings were retrieved from the Mariana Trench (M.289, station 225) at a depth of 4475 fathoms.

#### Dredge

Tizard et al. (1881) describe in detail the make-up of the dredge which was a weighted iron oblong framework with a muslin sack attached. The ship carried several sizes with smaller dredges used for deeper work up to 3875 fathoms (Station 25). Usually either the dredge or the trawl was employed but very occasionally both appear on the bottle labels at the same collecting station (eg. M.195, station 162). The dredge line was weighted further up the rope so that the dredge was dragged across the bottom layer when the coal fired engines onboard were started. Once at the surface, the contents of the bag were tipped onto the deck for viewing and sampling. It must be noted that the depth to which the dredge sampled below the water-sediment interface cannot be calculated, or the amount of sediment lost as the dredge was hauled to the surface. There are no documented details of the size of the mesh used. It’s clear from the narrative to the cruise that sometimes a great deal of sediment was retrieved so samples taken represent only a small fraction of dredge samples taken. Sometimes a smaller muslin bag was placed in the bottom of the dredge so that finer sediment could be retained if there was a possibility that it would be lost on the ascent to the surface.

#### Trawl

The trawl (Figure 3d) operated on a similar principle to the dredge but was designed differently as a weighted beam was attached to the sampling bag so that the beam agitated the surface sediments as the ship moved slowly drawing the trawl along the bottom. The deepest Trawl sampling is at 3000 fathoms at Station 264 but it was also used at very shallow depths.

**Figure 3.**
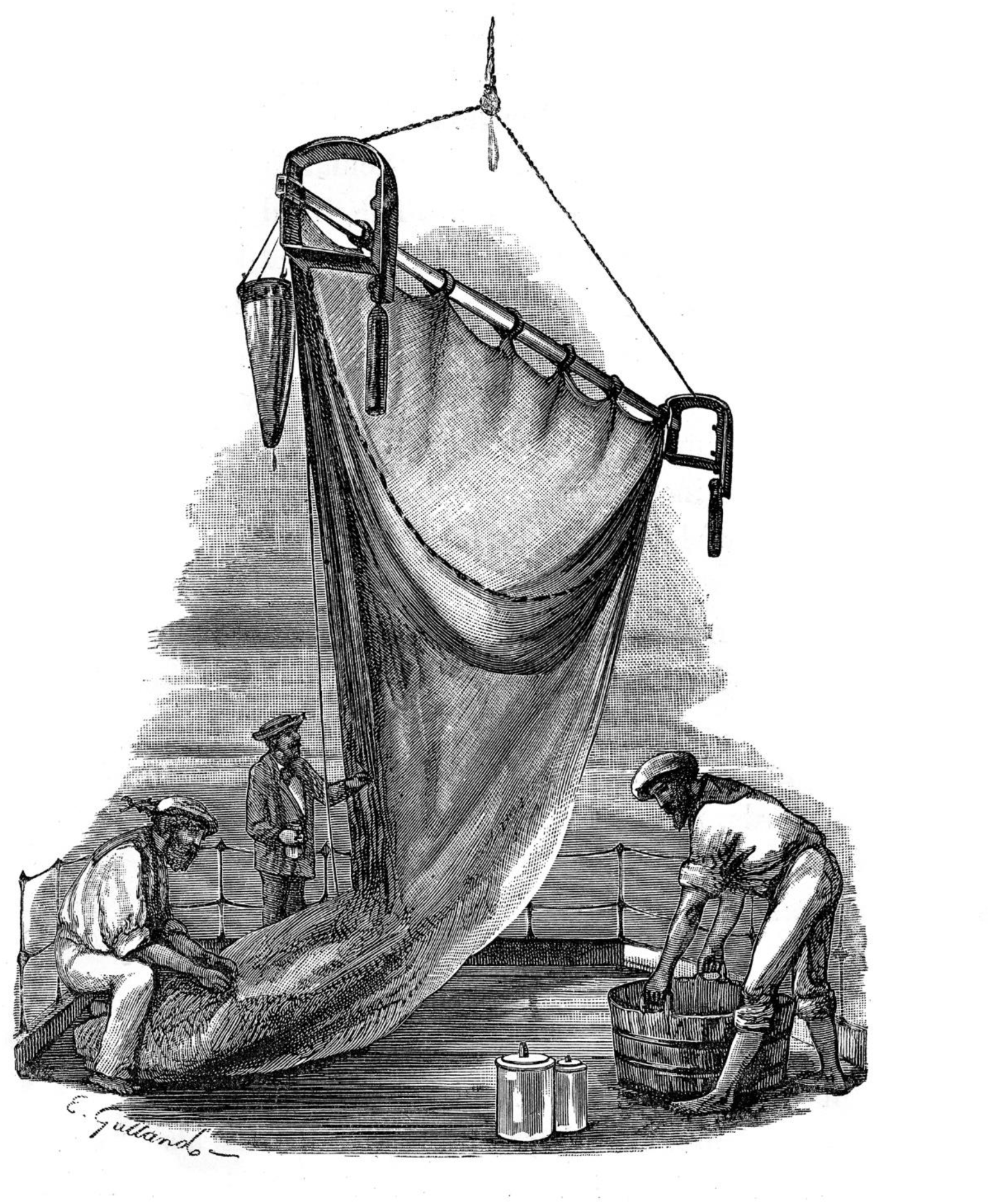
Tow net at dredge/trawl. (from Tizard et al. 1885)

#### Water Bottle

Water bottles were often attached to the sounding line to collect any suspended sediment from the water column. Samples M.65 and M.662(2) both from station 42, are the only instances where water bottles are specifically mentioned on labels but there are many references to bottle collections in Tizard et al. (1885).

#### Tow net, plankton

At all sampling locations and sometimes at points between named stations, collections were made at various depths using a plankton tow net (Figure 3e). These collections are often represented by Canada Balsam slides that indicate “surface” at 10, 20 and max 100 fathoms, although many slides just say “tow net”. The collection at the Natural History Museum of these slides does not cover the whole of the voyage and are often biased towards locations where planktonic foraminifera were well represented. “No systematic attempt was made to form complete collections of pelagic organisms

representing each region of the ocean visited by the expedition. To have preserved all of the collections from the surface waters would have been beyond the resources at our command” (Wyville Thomson, 1877, p. 1274). There are also tubes of “loose” material labelled “surface animals” that include larger items of Mesoplankton. At some points, particularly later in the cruise (in the collection it is after M.278, station 218), the crew experimented with connecting tow nets to the dredge, trawl and sometimes “at weights” (Wood, 2024) with the intention to collect suspended material that may have been dislodged from the surface when the trawl or dredge travelled along the bottom (Figure 3).

#### Anchor mud

Collections of anchor mud are present from Manila Harbour, Humboldt Bay Papua New Guinea, Nares Harbour PNG, Honolulu Harbour, Stanley Harbour Falklands, off Ascension Island and Vigo Bay, Spain. This is a small number of the anchorage sites so presumably not all provided sediment suitable for sampling.

#### From boat

There is only one reference on the labels to a sounding taken from a Gun Boat off Bermuda by Bethel (M.57). However, a smaller 30ft steam boat was employed for dredging and taking soundings in shallower water (Figure 4). Another boat related sampling was some sand from the arsenal of H.M.S. Challenger at Tahiti (M.348).

**Figure 4.**
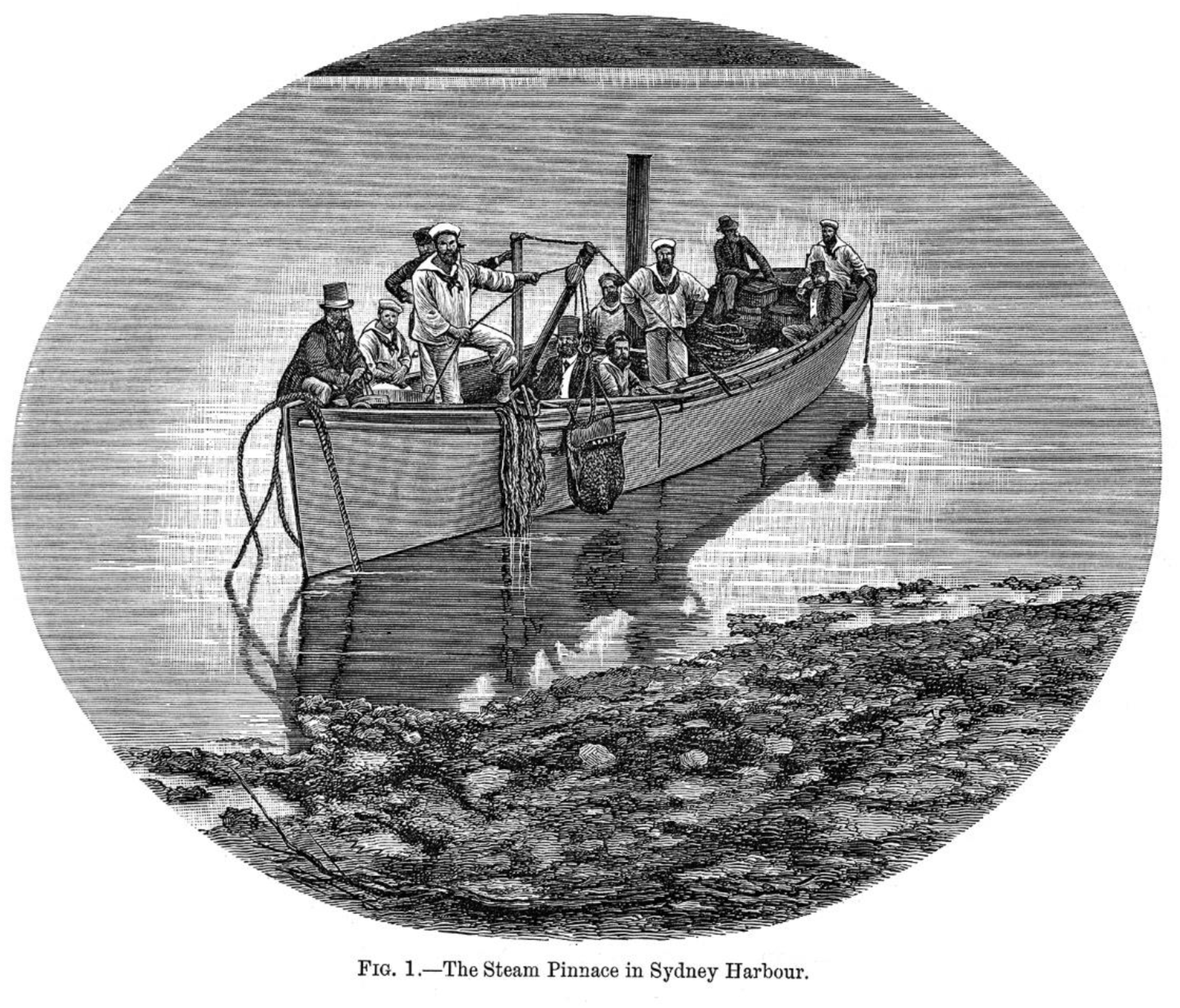
The Steam Pinnace employed for taking shallow water dredges (Tizard et al. 1885, fig. 1)

#### Collecting variations

Throughout the voyage and particularly at the start, variations were employed in the methods for collecting. Murray notes (Murray, 1895, p. 1273) that early on in the cruise, ‘small samples only of the deposits were preserved’ since they estimated that the quantity collected exceeded was larger than what was needed for any scientific purpose. This thinking was reviewed and in the later years of the voyage, much larger quantities were preserved and taken home. Towards the end of the cruise, the amount of sediment sampled appears to increase with M.408 at Station 338 represented by 32 rock bottles of sediment compared to most other stations where one rock bottle of dredge sediment was sampled (Fig. 5). However, Murray does also note that all deposit from the trawl or dredge was sieved and larger particles were obtained throughout the voyage. The Narrative (Tizard et al. 1885), Murray and Renard (1891) and the Summary (Murray, 1895) all indicate relative levels of success for the methods employed, for example, at Station 8 the dredge came up empty (Murray and Renard, 1891, p. 43) and at Station 9 the sounding tube penetrated over a foot into the sediment. Sometimes different methods eg trawl or dredge were employed because of the depth of sounding. A complete listing of sampling sites is included in Tizard (1885) Appendix 1. However, there are some inconsistencies between the methods listed there and the material present in the collection according to the labelling. For example, Tizard suggest that the method of collecting at Station 164B should be Trawl but the material in the collection is labelled as Dredge.

**Figure 5.**
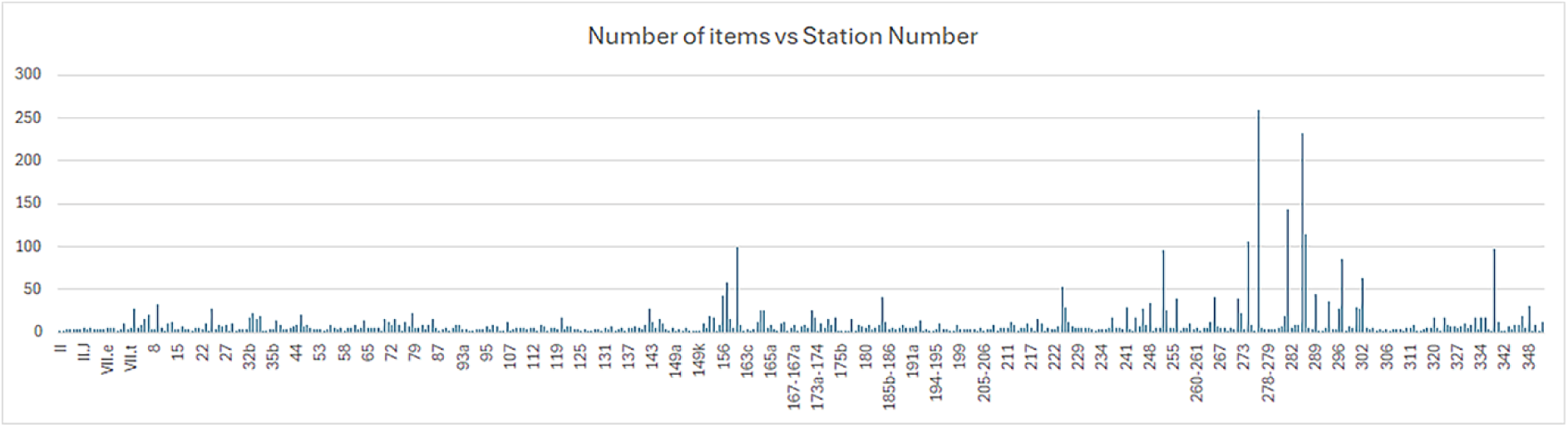
Number of collection items plotted against station number.

### From H.M.S. Challenger to the British Museum (Natural History)

Murray (1885) relates how no definite arrangements had been made to distribute the biological work amongst the naturalists on board and what was to be done with the collections that accumulated during the expedition. After a few months, there was an excessive amount of material in the workrooms and some of it began to suffer. The task of preserving, labelling, packing and storing the collections was enormous and although initially, some of the Naturalists would have liked to keep them all on board, it was agreed that they should be sent back at regular intervals to William Turner, Professor of Anatomy at Edinburgh University, who would examine their condition and take charge of them until the expedition’s return. Each Naturalist was responsible for one or more collection of specimens and it was John Murray who oversaw the Deep-Sea Deposits. None of them were allowed to send home private collections of their own.

Carefully packed collections were sent back to Edinburgh from six different ports: Halifax, Bermuda, Cape Town, Hong Kong and Sydney. The Royal Mail had an extensive network of mail ships which were reliable and links were also made with rail travel. Jones (2023) describes how 65 large boxes and 10 casks of natural history objects which departed Sydney on 6 June 1874 for San Francisco on a ship encountered difficulties in the Pacific but arrived safely 32 days later, where they were loaded on railway carriages bound for New York City. There, they were loaded onto a steamer bound for Britain and arrived safely in Edinburgh at the end of July 1874.

Once in Edinburgh, Professor Turner’s job was to unpack the collections, check the jars for breakages and refill them if necessary (Jones 2023). As leader of the scientific studies, it was Wyville Thomson’s responsibility to see this plan carried out. However, even he could not wait and he sent some

samples to eminent professors back in Britain before H.M.S. Challenger’s return (Lingwood, 1981). Moseley was also guilty of insatiable curiosity and Joseph Hooker, Head of Kew Gardens had already described the plant specimens he had been sent by the time the ship came home. Once the expedition was over, ‘the first official distribution’ (see Lingwood, 1981) was carried out in 1876 and the colossal task of writing the reports began. John Murray contributed to seven of the 50 Challenger Reports, including the Deep Sea Deposits report, with the Belgian Abbe Renard as co-author. Alphonse-Francois Renard, was a Reverend and Professor of Geology and Mineralogy at the University of Ghent (Henriet, 2010). Some labels in the collection bear annotations in French that were presumably made by him.

John Murray took over the leadership of the Challenger Report publishing in March 1882 after Wyville Thomson’s death (Tizard et al. 1885). During the writing of the reports which was completed in 1895, the collections were housed at The Challenger Expedition Commission’s Office at 32 Queen St, Edinburgh.

Wyville Thomson enlisted scientists of international repute to examine the collections and he devised a scheme which eventually and unintentionally led to the dissemination of the collections far and wide: each specialist had to split the collection in order to make two identical sets of specimens (for description and drawing), one set was to be sent to the British Museum. The third set contained the rest of the specimens for that collection. It was understood that both sets of duplicates should be returned to the Challenger Office once the reports were written but this did not always happen, and many ended up in private collections. On 6^th^ March 1895, John Murray presented the ‘duplicate’ Challenger collection specimens, one from each station, to the Department of Geology at The BM(NH). These are present as three drawers of small glass lidded cardboard boxes in the Murray Collection (Figure 6).

**Figure 6.**
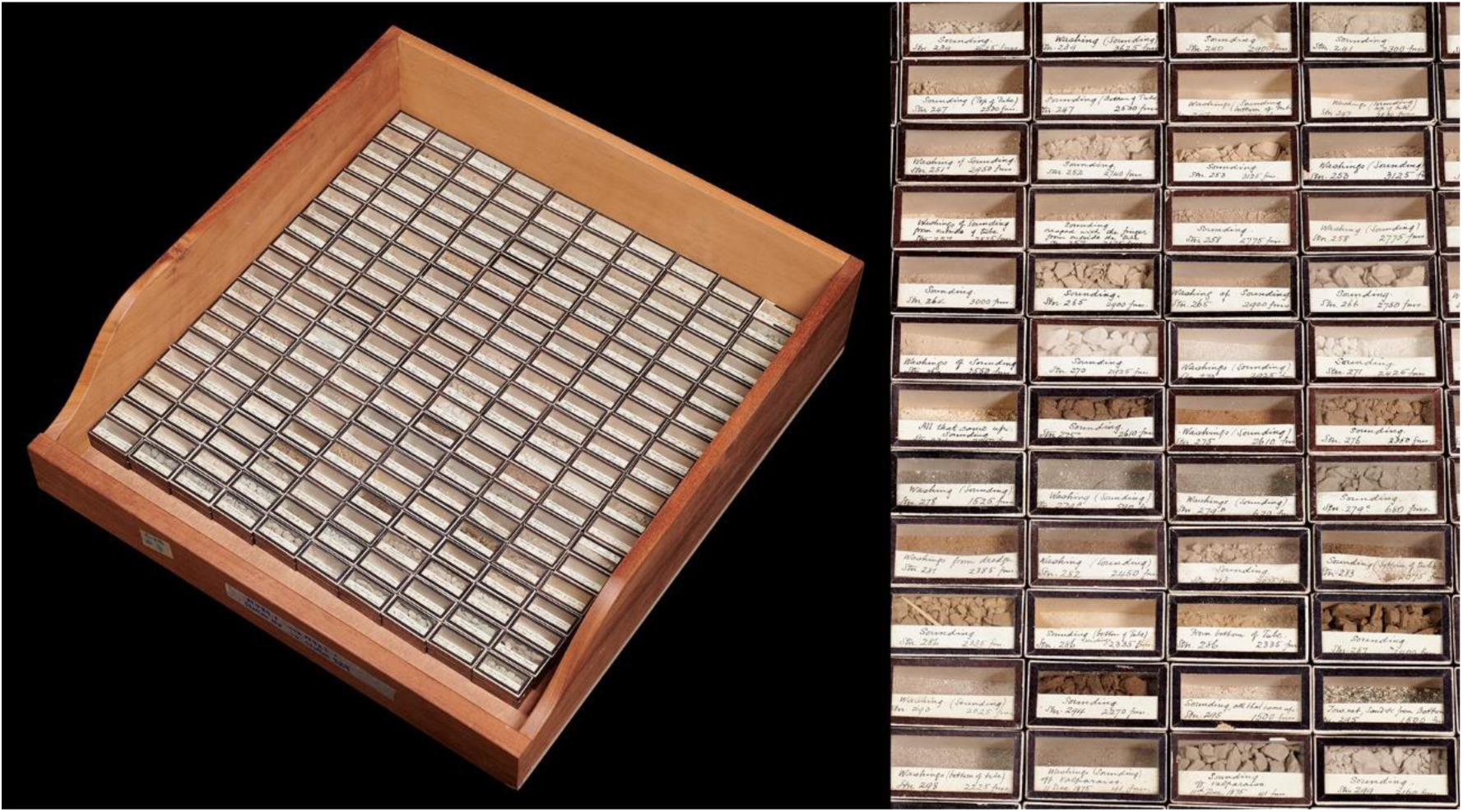
A drawer from the Duplicate Collection presented to the British Museum (Natural History) in 1895, one sample from each station.

Fittingly, for collections born out of a round the world voyage, they were soon on the move too: first to 45 Frederick St, Edinburgh in 1890 and then to Villa Medusa, 14 Boswall Road, Edinburgh, in 1904 where the whole of the Murray Sediment Collection was kept and curated. Sadly in 1914, John Murray died in a car accident. In 1919, following discussions with the Murray family about the fate of the Murray Collection, Edward Heron-Allen (see Hodgkinson and Whittaker, 2004), a prominent polymath, and his assistant, Arthur Earland of the BM(NH) found the soundings and dredgings of HMS Challenger in good condition, in the Deposits Room of Villa Medusa, occupying a total of 51.5m of shelving. They recommended that all the main collection of 9,746 Marine Deposits, soundings etc, together with the library, including the H.M.S. Challenger photographic plates and microscopic preparations should go to the BM(NH) to form the basis of the Oceanographical Collections (Kempe and Buckley, 1987).

### Collection materials and storage at the British Museum (Natural History)

Some clarification about the proper care and costs involved took place, involving the Murray family, the Trustees of the British Museum (Natural History) academics from Edinburgh and London. Finally, the John Murray Library and his sediment collection, to be known from then on as The Sir John Murray Collection, arrived in the Department of Zoology on the 13^th^ October 1921. From 1927 onwards, the collection was housed is what was known as Block B or the Discovery Hut (Kempe and Buckley, 1987, plate 9), still under the Department of Zoology, sharing the space with the Whale Research Unit of the ‘Discovery Investigations’. In November 1935, the Murray Collection was transferred to the Department of Mineralogy. In 1940, it was moved to the Museum’s basement, a below ground level bunker with thick concrete walls often referred to as ‘The War Room’ that now forms the nucleus of the museum’s Palaeontology Building that opened in 1977. This basement was flooded on 6^th^ March 1966 (see illustration in Kempe and Buckley, 1987, plate 12) but luckily only the labels on the outside of some of the jars were damaged. All Challenger Collection bottles that were affected have labels referring to the damage (see listing in Appendix 1.4).

To prepare for the new Palaeontology building, the oceanographic collections and research laboratories were moved in 1970 to temporary rented accommodation in North Acton and again in early 1980 to a larger outstation in Ruislip. The NHM purchased a site in London in the early 1990s and converted it to a collections store. This is the current resting place of the collection in eleven large double-doored mahogany cupboards by Waring and Gillow cabinet makers. A date of 1958 is written on some similar cabinets housing the later oceanographic collections but it is not clear if all of the cabinets are of that age and there is no evidence of flood damage from 1966. All manner of glass tubes, bottles, jars and boxes are arranged in chronological order of collecting from Station I to Station 364 (Figure 7).

**Figure 7.**
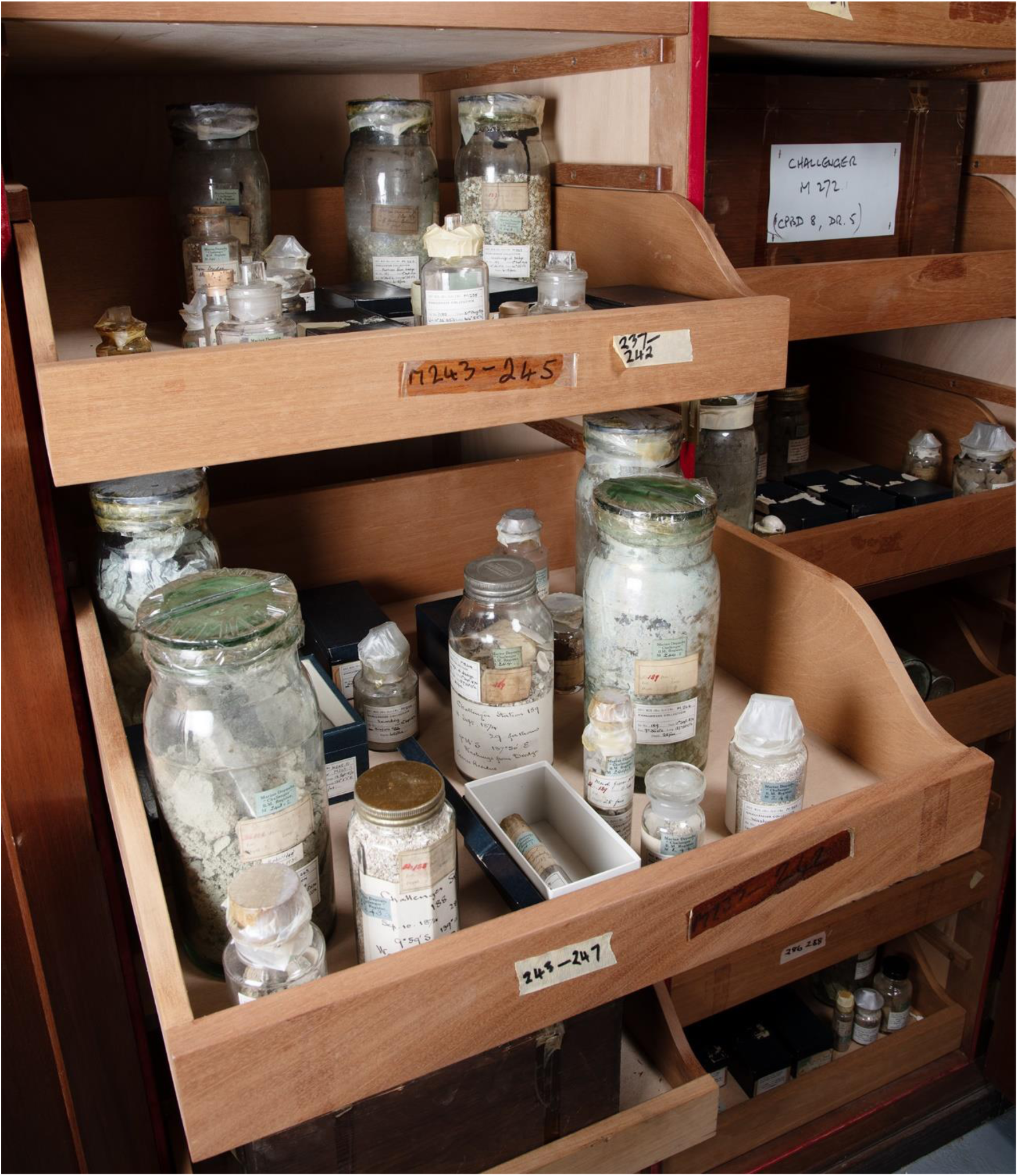
Current storage of the Sir John Murray Collection at the Natural History Museum.

#### Glassware

The largest bottles in the collection have been referred to as “rock bottles” (Figure 1a). The predominant variety has only a slightly developed neck and a loose lid that rests on a rim of cork. To attach these lids, a piece of animal skin was tied to the neck of the bottle by a piece of cord. The skin and cord, when present has then been painted with a black tar-like substance to seal the top, although there are some occasions for smaller bottles where the tar has not been applied or may have been removed. Only a small percentage of the rock/drop bottles (mainly shallow water deposits) have this original seal and two in the collection retain some original liquid, presumably spirit. From previous moves, many of the bottles are temporarily sealed with non-conservation grade materials including Sellotape and stretchy translucent material that has now become brittle. There is an enormous variety of diameters and sizes of bottles in the collection which makes consistent description difficult. For the purposes of the database we have described them first by their material (almost always glass for bottles), their approximate size (large, medium, small) and then their type of stopper or lid. These mainly have ground glass stoppers with a transverse bar to aid opening but small numbers have cork stoppers, plastic screw lids, metal screw lids or no lids at all.

#### Glass tubes (various sizes)

Sounding collections and sieved/analysed fractions are generally smaller and have been stored in a wide range of glass test tubes of varying sizes, each topped by a cork which mostly extends above the level of the top of the tube (Figure 8). Occasionally these corks have been inserted so they are flush with the top of the tube and are difficult to remove. Sets of glass tubes common to each collecting site have been stored within lidded cardboard boxes.

**Figure 8.**
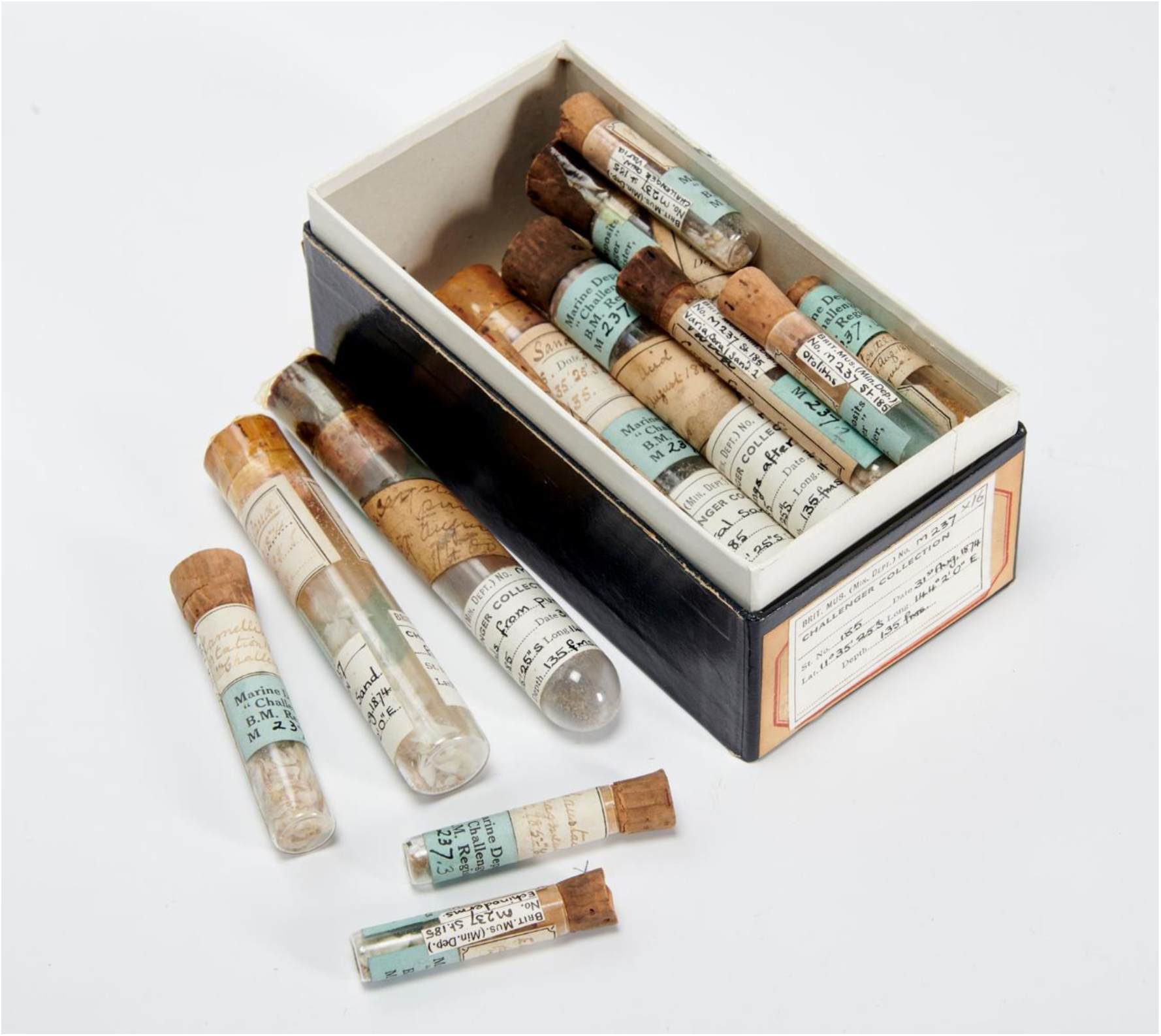
An assortment of glass tubes from NHMUK M.237.

It is difficult to tell whether all of the glassware is original. Annotations on labels in the collection suggest that rebottling of small parts of the collection took place in February 1956, August 1958,

March 1959, April 1959, May 1959, October 1959, November 1959 January 1960, August 1974, May 1975 and October 1993 (Appendix 1.1 for detailed list of rebottled material). Glassware variants (Figure 9) suggest that other sources of glassware may have been used as there are jars of Hookers Malted Milk (M.244(4)) and Horlicks (M.244(6)) which may or may not have been taken onboard the Challenger. There are a couple of examples of butter jars, one that was found broken on 1.10.1993 and has now been rebottled (M.339(10)). Another bears a raised marking around the rim of the glass top held on with a metal screw cap, reading “Masons Improved Butter May 10 1870” and the bottom is inscribed with “E364” (Figure 9) which suggests that food bottles and jars on board may have also been appropriated by the scientists. Another food reuse jar candidate mentions Branston Pickle (M.316(46)). There are also a small number of Kilner Jars present in the collection (eg M.256(7)). Another bottle is octagonal in section and bears the writing of the foraminiferal worker Arthur Earland (see later section on summary of work on the collection) so some bottles may have been added to the collection and presented back to John Murray after their contents had been studied. Other variants include a rock bottle with an “Aire and Calder, Castleford and London” glass stamp on the lid of the of jar (M.271(6)). Some rock bottles show an iridescence that represents devitrification/delamination of the glass rather than a chemical reaction with the bottle contents (Figure 1b). A figure illustrating some shorter bottles with pronounced lips that were presented to Nova Scotia Museum by the crew of the Challenger after their first Atlantic crossing (Davis 1973, fig. 3) are interesting in that they must have been amongst the first set of bottles loaded on board. Only a few of the type donated to Nova Scotia Museum are present in the current collection and these contain small notes to show that large bulk sediments available in large boxes but are stored out of sequence. It would suggest that John Murray or the other scientists on the Challenger may have used this type of glassware for other collections. A list of the bottle sizes and numbers purchased is available (Tizard et al. (1885) p.43) but an attempt to match all bottles in the current collection with the measurements was unsuccessful which would again suggest that there have been many changes to the glassware over time. Presumably stocks of glassware were also replenished at various points during the expedition. There is one example of a square section bottle and the flat tubes from the list of Tizard et al. (1885) are not present at all, although they are present in a box of residues exhibited as part of the museum touring Treasures exhibition (see later section on exhibits and National Museum of Nature and Science (2017), p. 8).

**Figure 9.**
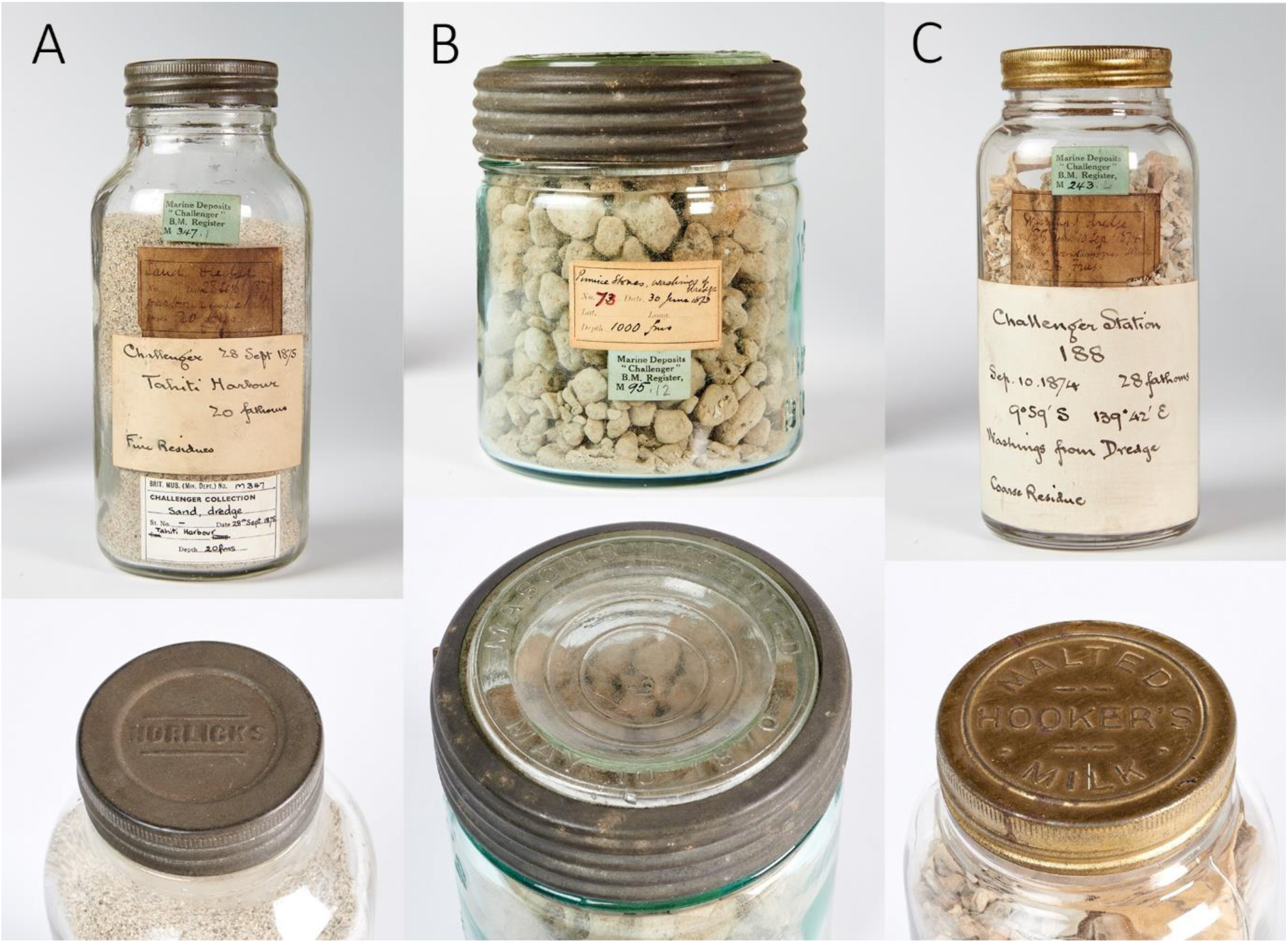
Variants and “re-used” glass bottles. A. Horlicks NHMUK M.347(1). B. Mason’s improved Butter jar with pumice, NHMUK M.95(2). C. Hooker’s Malted Milk NHMUK M.243(4).

#### Wooden boxes

144 Cigar boxes are listed as having been taken on the expedition (Tizard et al (1885), p. 43) but only a couple remain in the collection, albeit in a poor condition (Figure 10a). Many of these sub conservation grade boxes have been replaced by plastic or glass lidded boxes over the years. Large boxes containing many kilogrammes of sediment, pumice or nodules (Figure 10b) are also present but not listed as being taken. It would appear, due to the printed numbers that appear on some of these boxes, that they originally held collections of rock bottles. Wyville Thompson (1877, p. 16) recounts how these were kept in the boat magazine and their contents carefully logged so that the scientists would only need to ask a member of the crew to retrieve a specific box number if necessary.

**Figure 10.**
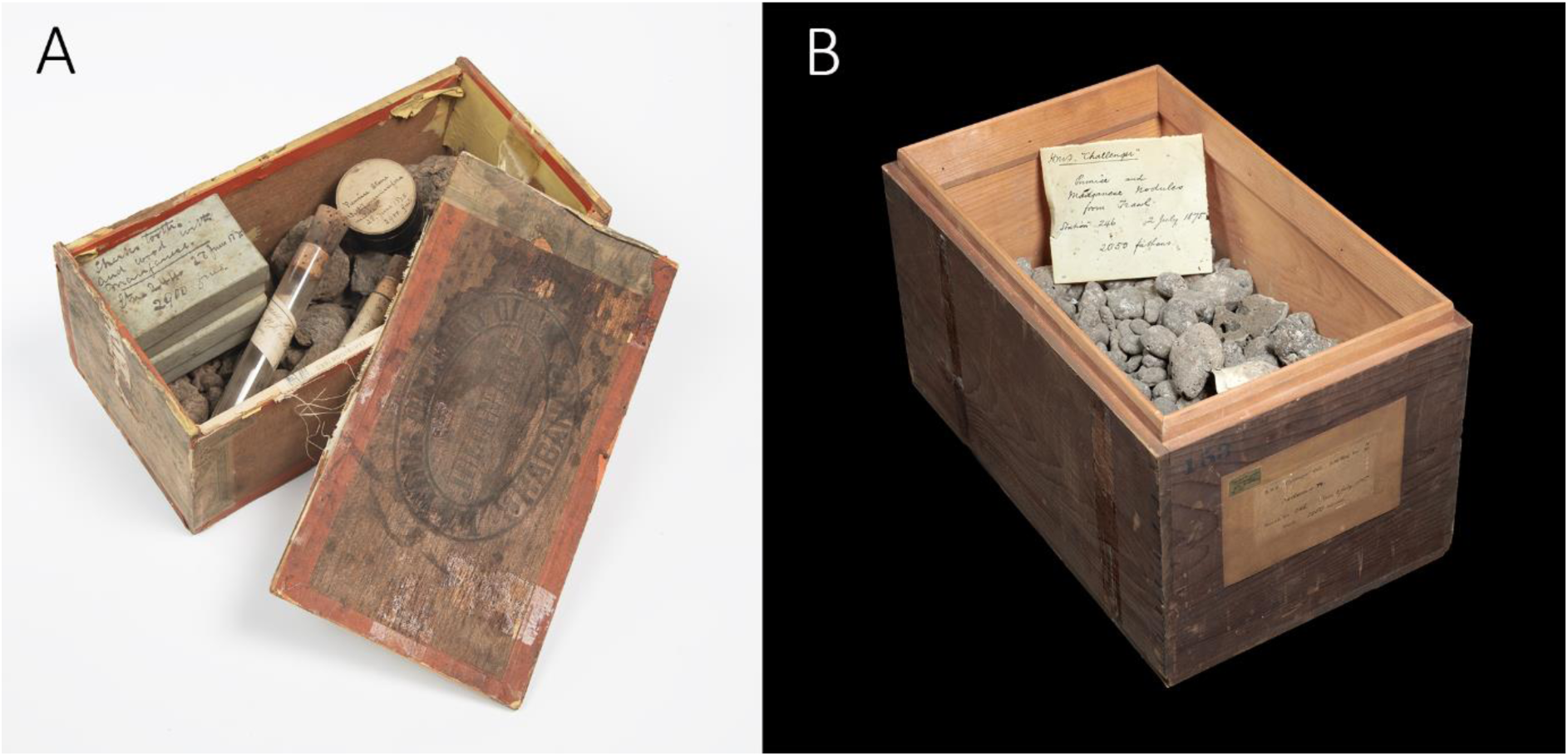
Wooden boxes. A. Habana cigar box containing, a pill box, two small cardboard boxes and glass tubes, NHMUK M.309(9). B. Large bulk sediment box with pumice NHMUK M.311(1).

#### Cardboard

There are various sizes of cardboard box with either a cardboard or glass lid. The cardboard lidded varieties are a faded bluey green and often have Murray’s writing on them (see also box in Figure 10a). The duplicate collection that arrived in 1891 (Figure 6) displayed each sample in a small oblong outline cardboard box with a glass lid that shows the deposit and part of the label. Some larger versions are also present in the collection, but these are common thoughout the collections at the Natural History Museum and cannot be proven to be original. Other variants on cardboard include pill boxes of various sizes, these, like the glass lidded boxes, have a black, shiny body with a lighter coloured card lid to enable labelling. These are certainly original as they display the same handwriting used for all labels in the collection (presumably John Murray’s) and the list in Tizard et al. (1885) suggest that 1584 pill boxes of various sizes were taken. However, there is no sign in the collection of the 144 oval boxes as all pill boxes present are circular in section.

#### Plastic boxes

Clearly a more recent addition, the collection, particularly where multiple specimens of nodules, pumice or sharks teeth are stored (Figure 12). The boxes are various sizes and sometimes plastic tubes are used to store glass tubes.

**Figure 11.**
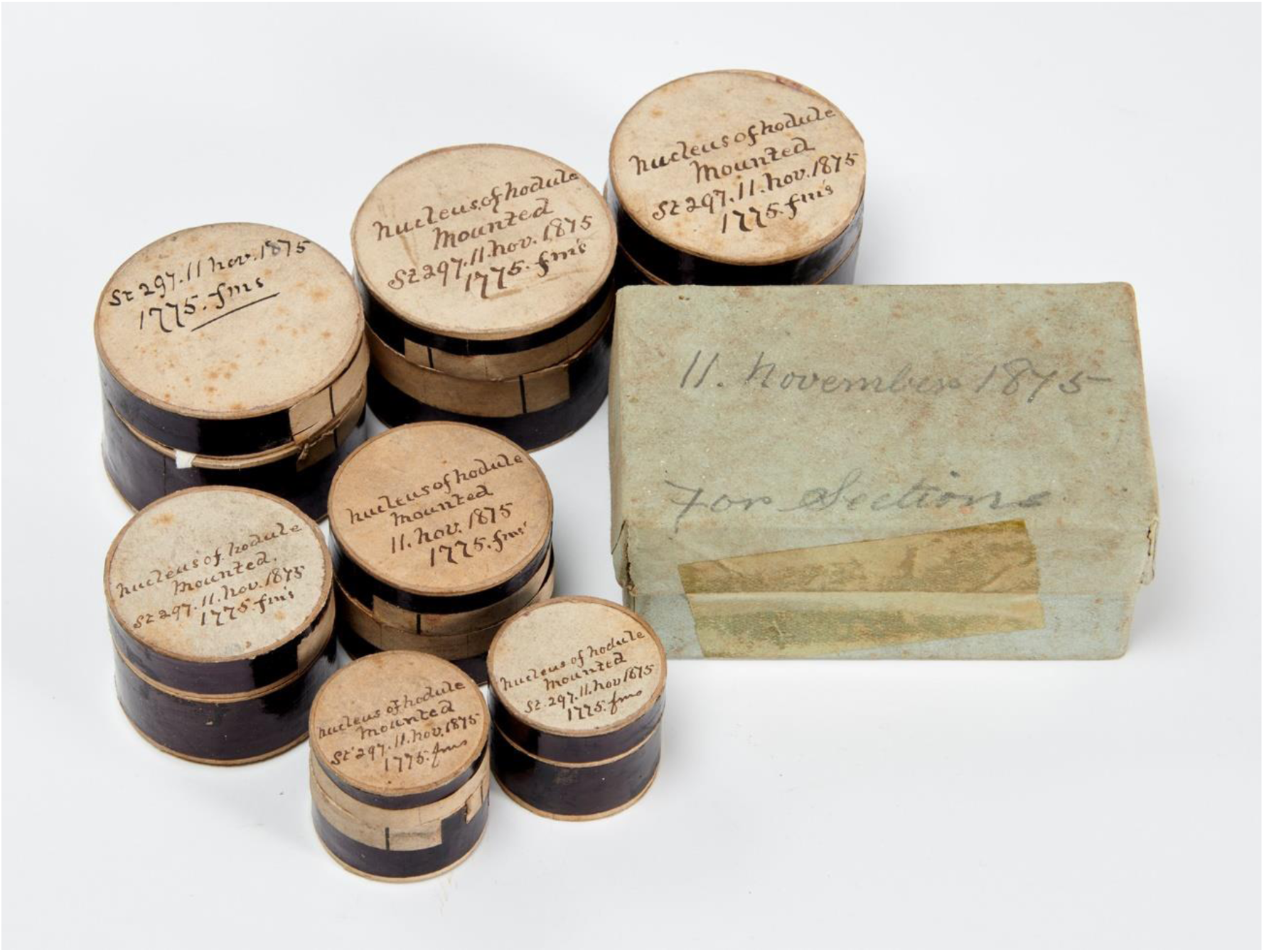
Cardboard pill boxes and a bluey green cardboard lidded box, NHMUK M.368(36)

**Figure 12.**
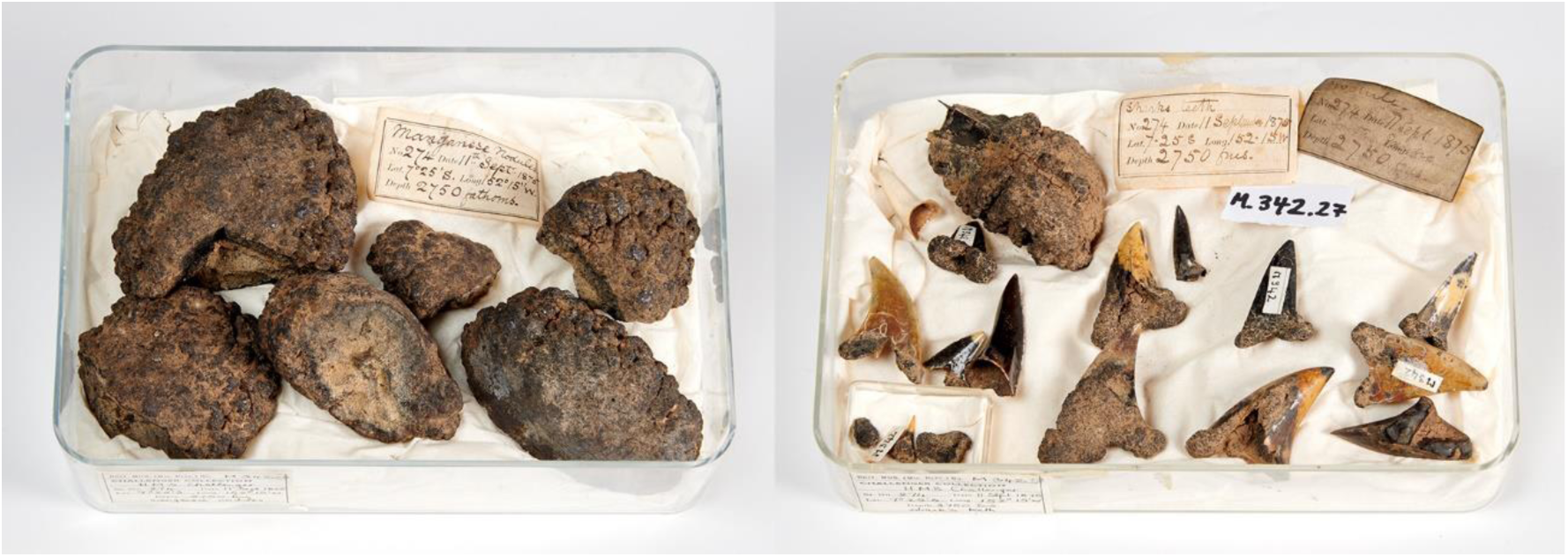
Plastic boxes with nodules (NHMUK M.342(28)) and sharks teeth (NHMUK M.342(27))

#### Figured material on boards

Specimens figured by Murray and Renard (1891, plates I-IX) aside from the thin sections, are present attached to wooden boards with yellow to brown printed exhibition labels attached to the upper surface of the boards (Figure 13). The specimens are held in place with steel pins that have sometimes been removed as contact with the specimens has potential to cause abrasion damage. The under-side of the boards often have original handwritten folded labels attached alongside more modern labels including the blue “M number” labels.

**Figure 13.**
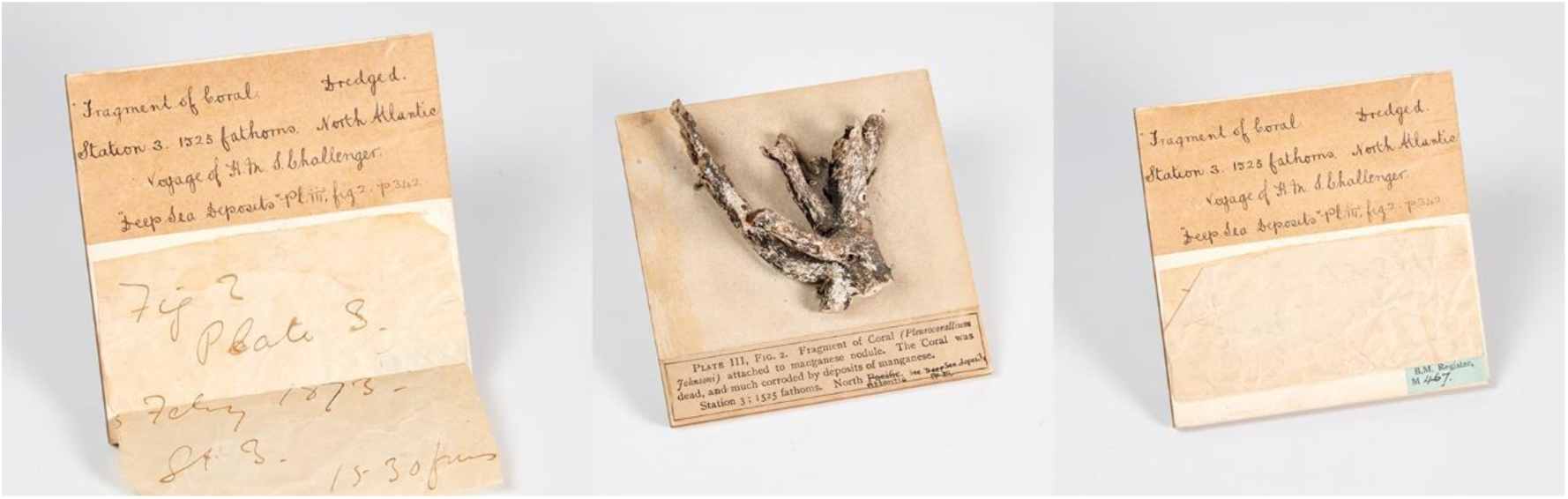
Figured material on an exhibition display board showing labels attached to under surface. A coral encrusted in Manganese from Station 3, plate III, fig. 2, Murray and Renard (1891), NHMUK M.467.

#### Slides

Slides are stored in a 105 drawer cabinet away from the main sediment collections at South Kensington. There are a wide variety of different slides (Figure 14) with sediment often embedded in resin and sectioned. Sometimes the sediments have been enclosed in circular glass cavities or wooden cavities topped with circular glass cover slips. Rather less successfully, some slides were constructed by sprinkling loose sediment on a thin layer of Canada Balsam and these residues have become loose in some slide drawers. Other slides have thin section preparations of nodules or pebble shaped igneous rocks, some of them almost twice as wide as a standard glass Natural History thin section preparation. Mostly these are too thick to enclose under a cover slip and like the open residues, these have sometimes come loose but are easily matched with the shapes in the balsam from whence they have come. Other preparations have also made their way into slide form particularly minerals including magnetic particles and residues from acidic preparations such as HCl or HNO_3_.

**Figure 14.**
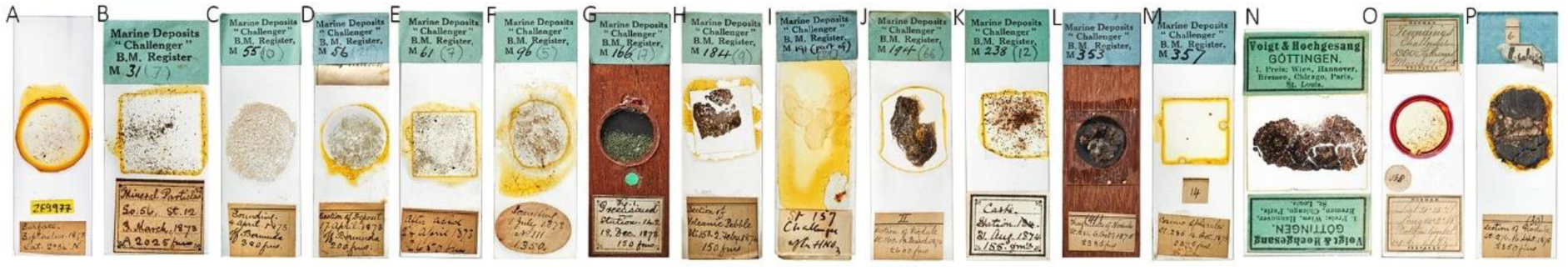
A variety of slides with specimen numbers all prefixed by NHMUK: A. plankton slide PM ZF 9977, B. mineral particles preparation M.31(7), C. open sounding strew, M.55(10), D. section of deposit mounted in copal, M.56(21), E. preparation after acid, M.61(7), F. sounding strew in Canada Balsam M.96(5), G. wooden cavity slide with loose sediment, M.166(17), H. rock thin section M.184(9), I. preparation after nitric acid and annotation by Ernst Haeckel M.191(29), J. section of nodule M.194(66), K. casts of Foraminifera M.238(12), L. wooden cavity slide with fragments of nodules M.353(xx) M. cosmic spherules in Canada Balsam M.357(136), N. Voigt and Hochgesang preparation of nodule, M.368(57), O. preparation of planktonic foraminifera by Norman M.408(65), P. open and thickly cut section of nodule M.344(169).

Some slides in the collection were made in Göttingen, Germany by Voigt and Hochgesang (eg M.208(23)) and another was prepared by the slide maker Norman (Figure 14). Other slides bear writing in French, possibly from the hand of Renard. A selection of 127 micro-preparations of mineral particles were belatedly returned to the British Museum on 22^nd^ November 1920 via Dr A. Schoep at the University of Ghent (written Gand on the descriptive typed note with the slides) and have registration number starting with that year. Presumably they were worked on by Renard in Ghent and retained for some reason by Dr A. Schoep for study. No publications appear to have directly related from this extended loan.

#### Alcohol preservation

All samples were originally preserved in spirit (presumably alcohol) on board (Wyville Thompson, 1877, p.16) but only a few samples with lids that are still sealed, retain any liquid (eg. Figure 1c, M.323(2)). Others include M.170(8), station 146 and M.417(5), station 346 while M.353(1) bears a note to say “in spirit until 1.4.1872” which is presumably 1972 as the sample was only collected in 1875. Original seals are relatively uncommon in the collection and have been seen in M.339(14) station 271, M.371(22) station 299, M.393(5) station 321, M.395(12) station 323, 26 of the rock bottles for M.408 station 338, M.417(8) station 346, M.418(3) station 347 and all of the M.421 rock bottles.

#### Other factors involved in preservation

For M.419(5) tow net at dredge and M.419(6) tow net at trawl, the original label reads “carefully dried as it came up but may have got a little coal dust in while drying”. Some of the associated plankton tow slides also include some black coal-like angular particles.

### Documentation

#### Labels and “Murray” M numbers

A very small number of original Challenger labels are so dark and discoloured that they are unreadable unless under uV light but have a standard shape and size with a black line around the margin (Figure 15). These have an initial blank line that allows a general description to be recorded and subsequently “No.” for station or sounding number, “Date” followed by “Lat.” and “Depth” on the bottom line. More modern labels have been applied that follow NHM labelling protocols including smaller blue labels that have a numbering system based on the prefix M denoting “Murray Collection”. This is a unique NHM sample prefix and was almost certainly applied after both parts of the Murray Challenger materials arrived at the museum as the “1895 Duplicate Collection” (Figure 6) have M numbers with a higher base number than the largely glassware-based main acquisition that arrived in 1919. Each M number generally reflects multiple items relating to a sampling site, although there are a few M numbers that cover details from more than one collecting site (eg M.120 = Stations 95 and 96) and sometimes a Station number can be represented by several M numbers (eg. Station 149B = M175-177). Some M numbers lack associated station numbers, particularly where they represent stops in nearshore locations and were sampled from less than 100 fathoms depth. The labelling of soundings have two numbering systems. The original labels use a sounding numbering system which is consistent with the preliminary report of Murray (1876), and this is sometimes recorded here on the ‘verbatim labels’. The ‘sounding’ number recorded in our dataset is interpreted from Appendix II of the Narrative of the Voyage (Tizard 1885, p. 1007-1015) based on the date and station number on the labels. These numbering systems do not match and it can be assumed the Tizard et al. (1885) numbers were allocated retrospectively when the collection had been returned to Edinburgh. However, we are confident that these refer to the same collecting sites even if the sounding numbers do not match.

**Figure 15.**
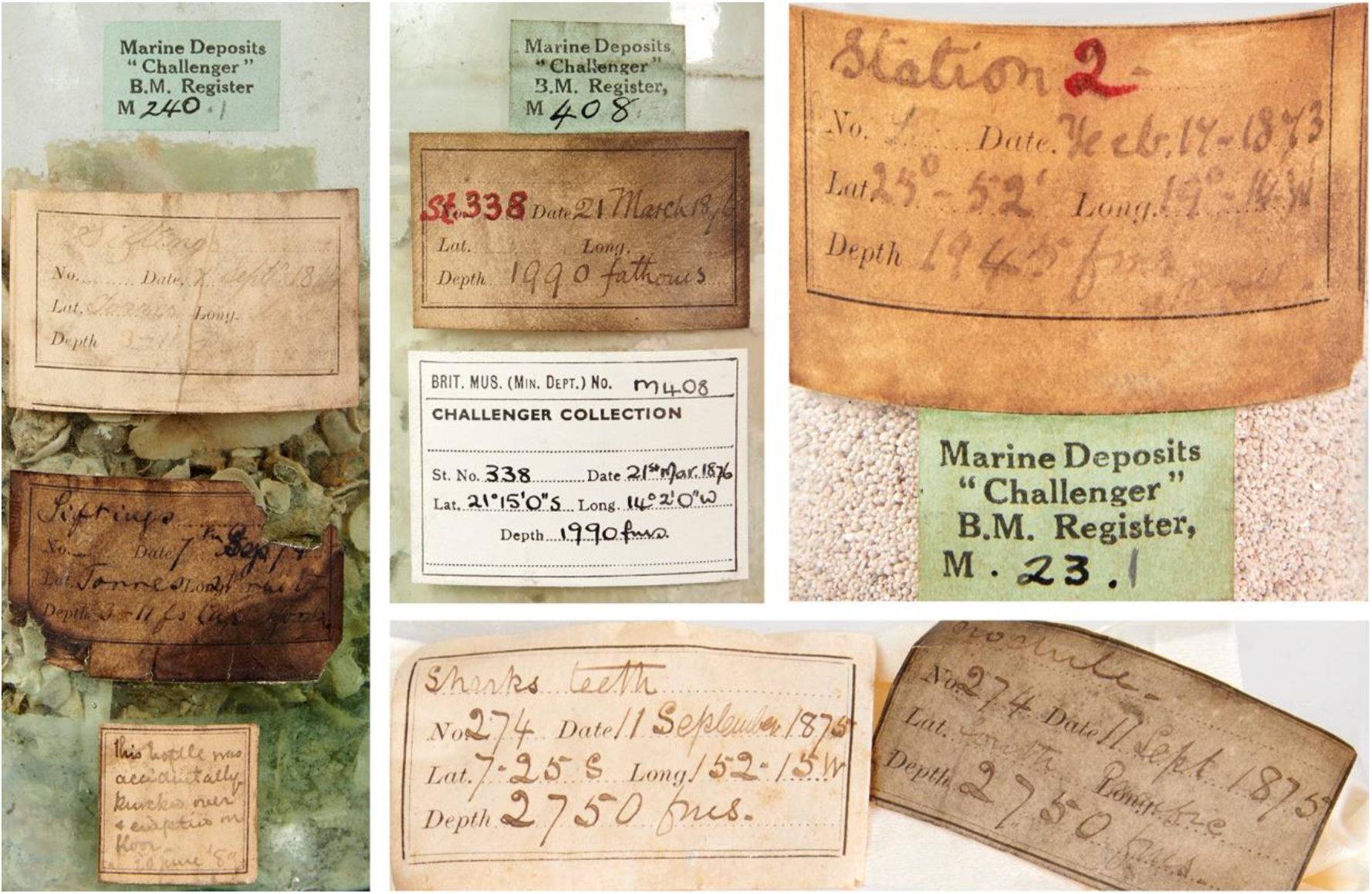
A variety of original and modern labels showing variation in preservation.

#### Inconsistent label data for collection method

A list of all sampled sites (Appendix II, Tizard et al. 1885) provides a details of collection method (Trawl or Dredge) and a numbered sounding record. There are some differences between this list and the handwritten labels on the collection. Sometimes the label states Dredge when Trawl is expected (eg. Station 184) and visa versa (eg. Station 194). Sometimes both are expected but only one present (eg. Station 232) where only Dredge material is present. Sometimes only one method is expected but both are present in the collection for example Station 195 where Trawl only is expected but both are present according to the labelling. Collection method is not recorded on some labels leaving it difficult to know if the material was derived from a sounding or dredge/trawl. In these instances the column has been left blank unless the Narrative (Tizard et al. 1885) indicates that only one collection type was possible or successful on that day.

#### Collections with no station number

A considerable number of collecting sites do not have associated Station Numbers and are not listed in Tizard et al. (1885). Georeferencable details of these are often given in the associated plans/maps and these interpreted latitude and longitude readings have been added to the dataset presented here after lat/longs have been interpreted via Google Earth (Table 1). Although some shallow water soundings less than 100 fathoms appear in Appendix II (Tizard et al. 1885), these unnumbered collections generally reflect samples taken at or near to sites where H.M.S. Challenger was moored in port and occasionally from on-land locations, for example beach sand.

**Table 1.**
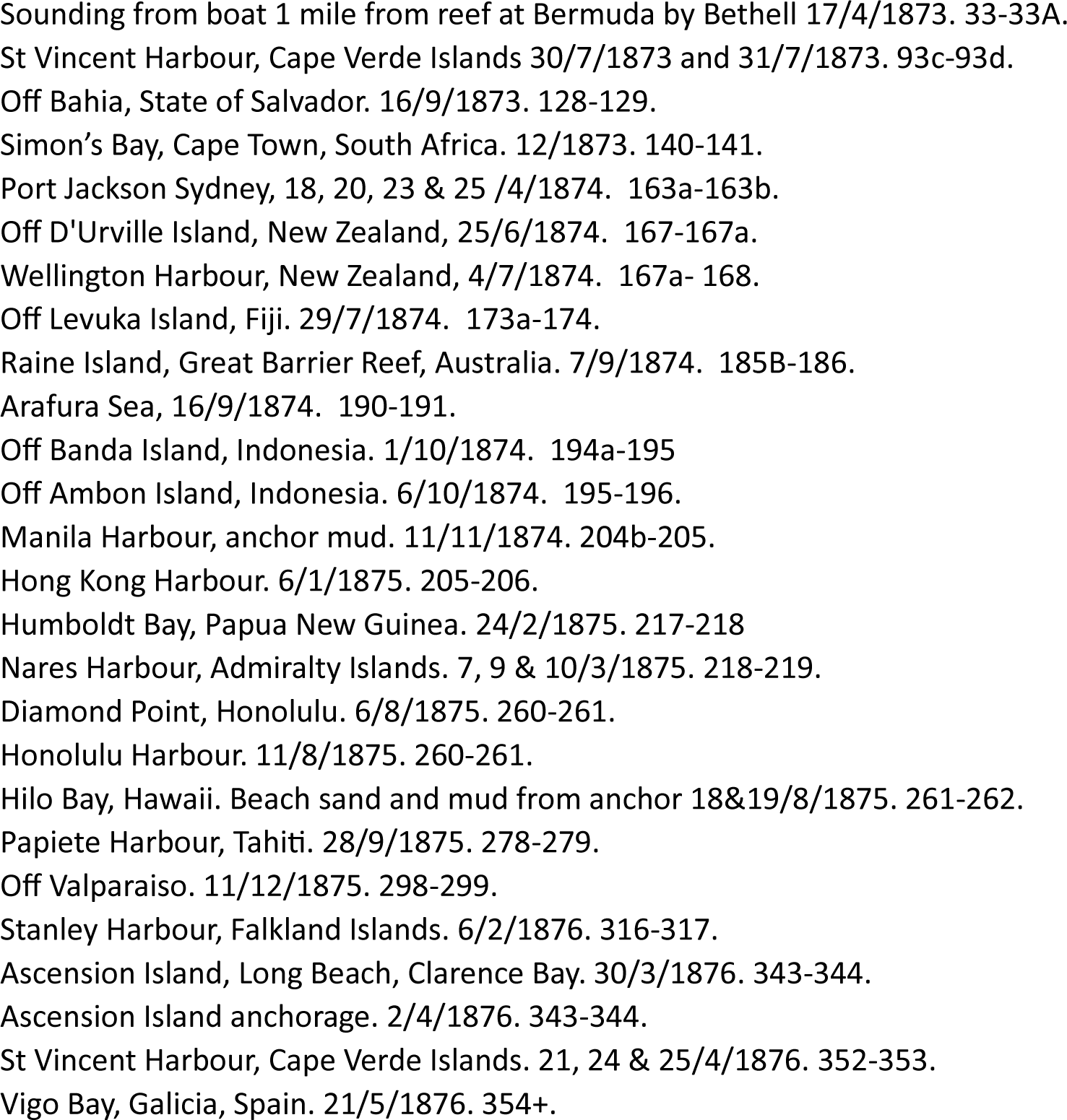
Additional non-station numbered sites: Brief geographical description, date of collection and Station number range.

#### Completeness of the collection according to available data

The collection does not have examples of material representing all of the sites listed by Tizard et al. (1885, Appendix II) but as mentioned above does have additional material collected from shallower sites near harbours and other nearshore points. Missing materials can sometimes be reconciled using details provided in the Narrative of the Cruise, for example M.31 from Station 12 states that the Dredge came up empty so the material in the collection most likely relates to the Sounding at that point. For other examples in the collection, it is not possible to do this as the tubes/jars/bottles give no indication of collection method for example there are two bottles and several tubes relating to M32, Station 13 that could relate to either the Sounding or Dredge taken at that point. M69 Station 47, the Narrative (Tizard et al. 1885) says that no Sounding was taken but the collection contains a sounding for that location.

Samples are absent from 99 numbered Stations (these are listed in Appendix 1.2) and this represents 19.8% of the total number of sites collected by HMS Challenger from named stations/sites according to Tizard et al. (1885). There are several explanations for the absent material. Many of these sites relate to the early part of the cruise where training for sounding and dredging took place and this was not always successful (eg stations III and IV). As the cruise went on, techniques were honed and the crew became more skilled at gathering samples. Often samples were not kept if previous closely located samples were very similar but usually, if the sounding tube only had a trace of mud on the outside or there was insufficient sediment recovered for analysis to have taken place. Other reasons were if the sounding was shallow or if the nature of the bottom was marked as ‘Hardground’ or only rocks were recovered. Taking into account the details given in the Deep Sea Deposits Report (Murray and Renard, 1891, p. 34-147) the number of expected samples that are absent reduces drastically to only 17 which is 3.4%. It is hoped that by publishing this dataset, we can identify if custodians of Challenger Sediments have collections that fill these gaps. The dataset also allows custodians to assess the relative uniqueness of their samples.

### Guide to the published dataset

The dataset in the form of an Excel file with three tabs is published at https://doi.org/10.5519/50h9e977 with descriptions of each column presented with other appendices relating to this paper (Appendix 1.3). The first tab is a listing of items in chronological order of collection and the second includes details of collecting sites with the two tabs linked by the site number from the first column on the second tab. This has been necessary as there is not a one to one relationship between Murray (M) numbers and collection sites. Some M numbers cover material from two different collecting events and sometimes a collecting event can be relevant to two different M numbers. Descriptions of all columns on each tab of the dataset are included in the appendix to this paper. The first two tabs are connected by Station Number. Where stations have not been allocated, a range of stations has been given. Verbatim label data is provided and interpreted information or comments on ambiguous data added as a note. All electronic records are present on the NHM’s data portal. A programme of digitisation over the following years will add images to each item in the collection and these will also be available via the portal.

#### Types of deposit present

Full details of descriptions, types and distributions of deposit are provided by Murray and Renard (1891). These are essentially grouped into terrigenous and pelagic. Terrigenous include “schistose rocks, shales, marls, greensands, chalks, phosphatic fragments, limestones, volcanic grits, quartzites, and sandstones” (p. xxviii, Murray and Renard 1891). Pelagic sediments from deeper water include blue mud, green mud, Globigerina ooze, red clay, pteropod ooze and diatom ooze. Some jars and boxes contain larger items for example, coral, bryozoans, sponges or other biogenic materials such as sharks teeth and whale ear bones. Some jars contain complete or fragmentary benthic molluscan material (Figure 1a). Polymetallic nodules or nodules of pumice are common in some parts of the collection, particularly from Pacific waters.

### Scientific use cases

The cases mentioned here are mainly derived from details gathered from labelling associated with the collection where details of past usage are recorded, usually scientists visiting or requesting loans from the collection to gather more comparative material to check findings published in the volumes. Many of these appear to be unpublished or difficult to connect with published records. It is important to note that this section is not intended to be a full coverage of past studies on materials obtained and published in the volumes, for example a review of studies on Challenger Foraminifera (eg Jones 1994). This section provides details where stakeholders returned to the original Murray sample collection for various studies.

#### Micrometeorites

Murray (1876) was the first to recognise that microscopic spherical objects in the sediments are of cosmic origin. Murray and Renard (1891, plate XXIII) figure some of these items that are currently housed with the meteorite collections at the Natural History Museum. Since then, Jedwab (1975) mentioned microscopic metallic flakes of iron, iron-nickel and nickel-iron from Stations 160, 276 and 286 and considered them cosmic in origin. A note with M.358(24) = station 286 says, “2 nodules sent to Prof J Jedwab for examination of cosmic spherules.” Former mineralogy PhD student Paul Yates analysed material from M.342(11), station 274 and M.358(12), station 286 with a note saying that it was returned in 1988. However, nothing was mentioned or figured in his thesis (Yates, 1993) in relation to these samples. More recently Hughes et al. (2023) report that micrometeorites have been obtained from M.194(49.1 and 50.1), M.316(50.1), M.333(24.1), M.342(16 and 50.1), M.344(53.1), M.357, M.358(1.1), M.408(40.1) using a combination of sieving, magnetic separation and CT scanning. These were Iron-dominated micrometeorites (or I-type micrometeorites) that have potential for us to better understand the storage and alteration of micrometeorites on the ocean floor by comparison with both pristine Antarctic micrometeorites and others recovered from the fossil record.

#### Ostracods

An entertaining label with M.350(1) reads, ‘Brady says - contains absolutely nothing’. It’s important to note that the brothers George Stewardson and Henry Bowman Brady were instrumental in describing the Challenger Ostracoda and Foraminifera with George Stewardson studying the Ostracoda (Brady, 1880; Davis and Horne, 1985) and Henry Bowman the Foraminifera (Brady, 1884). The Micropalaeontological Society’s highest medal awarded annually is named after them (Siveter, 2008). It’s not therefore clear which Brady the note is referring to. Samples M352(5) and M.408(16) both have labels: ‘Originally in Brady Colln. Trans from Zoo. Dept. March 1939’. Again, because it is not clear which Brady brother it relates to, this could equally be from the Foraminiferal collection which was stored with the zoological collections at the museum at that time (it is now with the Earth Science collections). Puri and Hulings (1976) designated some lectotypes of ostracod species originally described from the Challenger Expedition by Brady. Sample numbers M.198(5), M.220(7) and M.237(18) have annotations ‘Ostracods removed by H. Puri, 20.7.1967’ so it appears he revisited some of the original samples to gather additional specimens.

#### Foraminifera

Sample M.158(5) from Station 135c at Tristan da Cunha Island, Southeast Atlantic has a note with it that says “material originally borrowed by Prof. Bronnimann” but there is no date. Paul Bronnimann published extensively on benthic Foraminifera so it is difficult, without further details, to attach an outcome to this enquiry. Likewise, sample M.377(7) from Station 305b, Gulf of Penas, Chile has a label inside the tube saying ‘1973 subsample by R. Hodgkinson, BMNH, Pal Dep. (Protozoa section)’. Former Curator of Foraminifera at the Natural History Museum, Richard Hodgkinson, mentioned Challenger Foraminifera in his paper on the Pitcairn Islands (Whittaker and Hodgkinson, 1995) but it is not possible to link this request to that publication other than some of the taxa were originally described by Brady (1882). Sample M.414(1) also has a note inside the tube, ‘Returned by Lois McCormick July 1985, after *Borelis* investigation’. Lois McCormick was a curatorial assistant in what was the then Protozoa Section of the Palaeontology Department where the Recent and fossil Foraminifera collections were housed. It is clear that McCormick did not publish on this material but just over 10 years later, John Whittaker, then Head of the Micropalaeontology Division at the Museum, was a co-author on a paper on the foraminiferal genus *Borelis* (Jones et al., 2006) that includes reference to Brady’s monograph (Brady 1884) as well as Bob Jones’s Book (Jones, 1994) redescribing the Challenger Foraminifera. Another Natural History Museum staff member that regularly sampled the collection was Henry Buckley, former curator of the Ocean Bottom Deposit Collection from 1961 to 2001. Rillo et al. (2017) provide a detailed summary of Buckley’s work that included creating a taxonomic reference collection of planktonic foraminifera that he had processed and picked, partly from the Murray Challenger Sediment Collection but mainly from the museum’s vast Ocean Bottom Deposit Collection (Kempe and Buckley, 1987). Rillo et al. (2017) provide details of this collection via a dataset on the museum’s data portal (https://doi.org/10.5519/0035055). Marina Rillo imaged and registered the collection as part of her PhD project entitled *Unravelling macroecological patterns in extant planktonic Foraminifera* (Rillo 2019). Rillo also used specimens of planktonic foraminifera reprocessed from four original Murray collection sediments (M. 25, 192, 284 and 408). She compared the planktonic foraminiferal content of these samples with other data sets particularly the Holocene (last 11,700 years) ForCenS dataset from core top planktonic foraminiferal data derived from cores drilled since 1945 (Siccha and Kucera, 2017) and a dataset representing the Last Glacial Maximum 21,000 years ago (subsequently presented by Jonkers et al. 2024). The conclusions of these studies were that planktonic foraminiferal content of sediments can be used to investigate whether they are a true representation of the oceanic conditions of 150 years ago, suggesting that museum collections such as these have potential to act as a 19^th^ Century baseline for oceanic conditions. Rillo et al. (2020) also used these four Challenger samples as part of a study to evaluate the drivers of intraspecific size variations and to investigate any biases that Henry Buckley may have introduced while compiling the slides in his collection. The main conclusions were that Buckley did a good job in picking out all the key taxa but that there may have been a bias towards larger and more easily identifiable specimens. Rillo’s imaged specimens were amalgamated with a dataset from Yale University to compile a larger dataset of planktonic foraminifera that have been used as an identification training dataset (https://endlessforams.org/) and as part of a convoluted neural network to automatically identify planktonic foraminiferal species (Hsiang et al. 2019). This dataset has been used by several teams to investigate the possibility of using AI to quickly identify specimens of planktonic foraminifera with the objective of quickly building up large datasets of species occurrences. Fox et al. (2020) studied tow net samples from M.340, station 272 and compared the planktonic foraminiferal shell thickness with specimens collected almost 140 years later as part of the Tara Oceans expedition (Pesant et al. 2015). CT scans showed that the wall thickness was considerably reduced in modern forms with the conclusion being that this was evidence of the actions of ocean acidification (Fox et al. 2020; Vaccaro 2019). More recently the sediment collection and particularly some of the sieved residues have been used in unpublished masters projects investigating factors influencing the biodiversity of planktonic foraminifera (eg Satija, 2024). Zarkogiannis et al. (2025) used X-ray micro-computed tomography to scan small vials of Challenger sediment from the Murray Collection marked as plankton tow net and showed that the foraminiferal content can indicate whether benthic material is also present. They also demonstrated that there were two types of plankton tow net material collected by H.M.S. Challenger, nets attached to the dredge, trawl or weights and those trawled from the top 100 fathoms of the water column.

#### Diatoms and Radiolarians

A label with sample M28(14) shows that “4g of this spec. sent to Dr Riedel of California, 1959”. Riedel had published a paper on the sediment from station 225, describing the diatom *Ethmodiscus rex* (Rattray) and suggesting that the sediment was Middle Tertiary in age (Riedel 1953). Sample M344(1) was “Received from Botany Dpt Oct 1962. Contains no diatoms.” It should be noted that the diatom collections at the museum are still housed with the Botanical collections. Challenger diatom samples, part of the Comber Collection are present from stations 298, Hong Kong Harbour (between stations 205-206) and a raw sample labelled as Antarctic surface (Comber 442). Some Murray collection material is also present (M.347 from Tahiti Harbour).

#### Benthic mollusca

A label in the collection with M.324(4) from station 260 off Honolulu says “30cm^3^ sorted by Alison Kay to be compared with modern material”. Kay, who was then at the University of Hawaii, Honolulu, redescribed (Kay 1965) the Marine Mollusca from the Cuming Collection at the British Museum (Natural History). Although there are several references in synonymy lists to the work of Watson (1886), who described the Scaphopoda and Gastropoda from Challenger, there are no indications that Challenger material was directly used.

#### Polymetallic nodules

Deep sea manganese nodules were first discovered on 18 February 1873, 300 km south-west of the island of Ferro in the Canary Group (Station 3) (Murray and Renard 1891). Manganese nodules are often termed polymetallic nodules as several key metals can be recovered from them including Nickel, Cobalt and Copper as well as rare earth elements (see Glasby, 1977 and 2013 for a summary on their importance). Notes with the collection point to many requests for material to analyse including that 5.25g of manganese nodule powder was given to Prof H.W. Menard, California in Oct. 1959 from the first cited occurrence (M.24(4), Station 3). Many of these analysed materials do not seem to have made their way into the literature. There are also annotations in French presumably from Renard for example M344(76) ‘il fut impossible d’analyser ces centres de nodule’. [It was impossible to analyse the centres of these nodules]. A note on blue paper with M.372(12) from Station 300 says: ‘3 pieces of tufa coated with manganese from this box given to Mr Anderson, 21.4.1893’. A pink note says ‘3Mn nodules to Dr Kemp, 23.1.76’. An undated note with M361(37) Station 289 says “1/2 Mn nodule to D.S. Cronan, Royal School of Mines”. Another undated note suggests 10.8g of M25(30) from Station 3 was sent to Dr Riley, Liverpool Uni but presumably went towards the annotated record of detailed examination of Mn nodules from the Challenger Expedition (Riley and Sinhaseni, 1958). An interesting addition to the acknowledgements for this paper says “The authors wish to thank the trustees of the British Museum (Natural History) for the gift of nodules taken at Challenger Stations, 248 and 252.” More recently Dekov et al. (2010) presented geochemical details of metallic sediments from 6 samples between stations 292 and 302 and Monget (2016) a dataset of all manganese nodule occurrences in the the Murray Collection.

#### Other rock analyses

Pelagonite, a water altered volcanic glass from M373(24-30), Station 302 was sent to Renard in Feb 1879 and the largest and best ones on 28^th^ Dec 1879.

#### Sharks

Teeth of the extinct shark *Otodus megalodon* illustrated from the Pacific Ocean by Murray and Renard (1891, pl. 5 figs 1 and 2) have been involved in the debate over whether this shark is extant, although there have been no suggestions that the Challenger material directly supports this theory. Tschernezky (1959) used their maximum accumulated surface thickness of manganese dioxide (MnO_2_) and an assumed accumulation rate to estimate that M.482 is 24,267 years old (late Pleistocene) and M.481 is 11,333 years old (early Holocene). Greenfield (2023, fig. 1) re-illustrated these two specimens (Figure 16) and provides an interesting critical summary of these studies, concluding that, “there is no compelling evidence for extant *O. megalodon* and ample proof of its extinction” (Greenfield, 2023, p. 332). In 2024, based on unpublished data from the dataset presented here, new samples were made for dating analysis of *Megalodon* specimens collected by H.M.S. Challenger from the Pacific and Indian Oceans.

**Figure 16.**
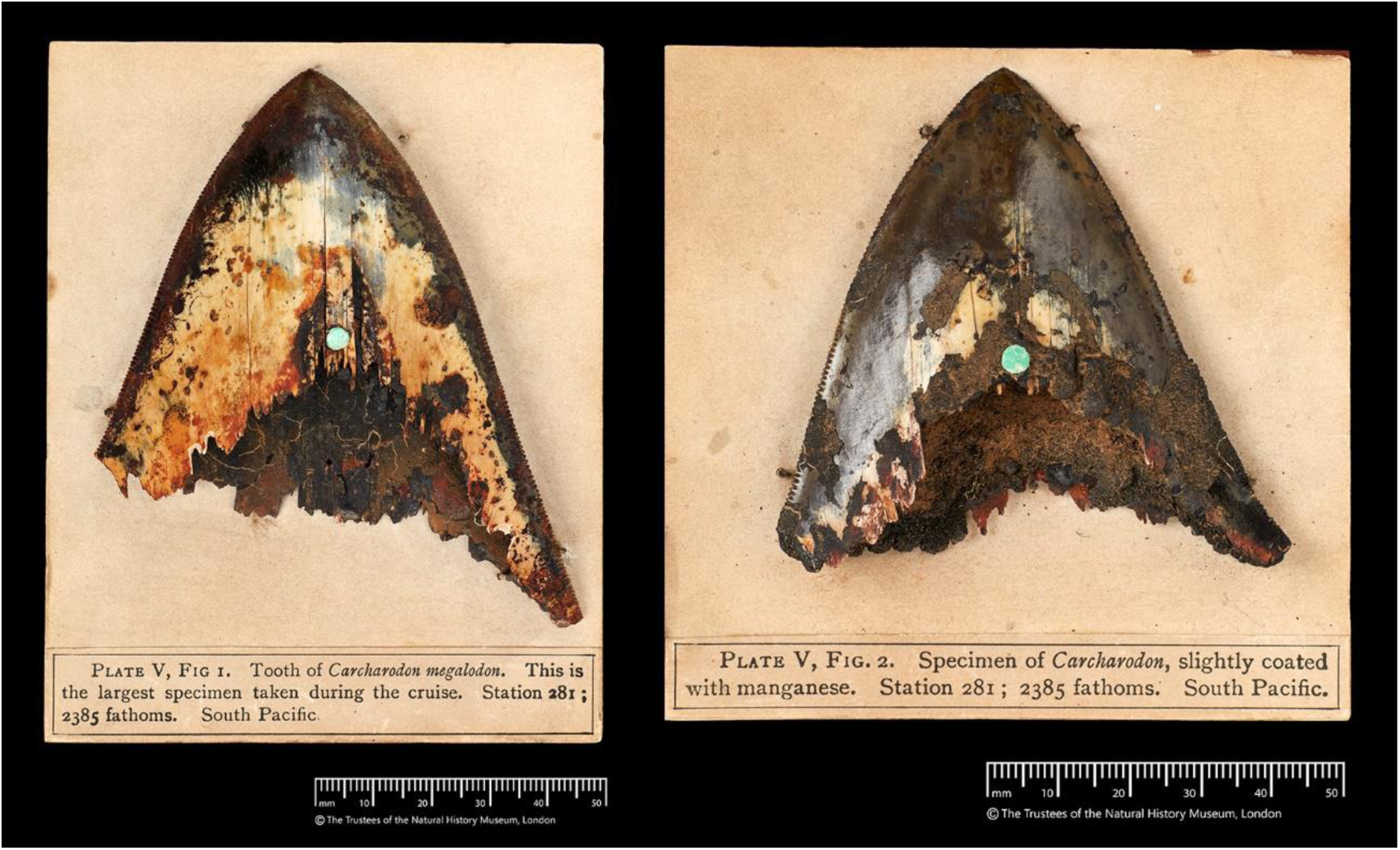
2 Shark teeth specimens on exhibition display boards (NHMUK M.481 and M.482)

#### Whales

Specimens of Cetacea from three plates from Murray and Renard (1891, pls VII-VIII, X; M.505-517, M.358(38-42)) have been found within this collection and include bullae, a brain case as well as tympanic, typano-perioitic, petrus and mesorostral bones. Another set of items (M.551-555) figured in the Cetacean volume (Turner, 1880, plate II) are mainly ear bones from ziphioids. A note with M.358(19) from Station 286 says, “lge piece of whale bone, spec. no.1 LOAN TO N. Higgs 6.3.2010. Zoology, NHM.” It’s clear that this hasn’t been published but it should also be noted that the whole collection of whale related items including unfigured specimens, was transferred to the Life Sciences Mammals Section in August 2024.

#### Other studies

A suite of analysed samples in the collection and marked “Analysis 1-9” covers earbones, manganese nodules, sharks teeth, nodules with sharks teeth, nodules with nuclei and a red cherty layer. These were taken from M.357 Station 285 but it is not clear who made the analyses and what sort of analysis was carried out as details remain unpublished. Two small B&W photographs of sections taken by Dr W. Mackie in 1910 are present in a cardboard box containing samples M377.1-M377.4 from station 305a,b. It’s interesting to note that there are no slides present in the collection that would have been the origin of these images.

### Exhibitions

#### Semi-permanent display

There is currently a small glass bottle of material from the deepest sample taken (M289.2, Station 225, 4475 fathoms) from the Mariana Trench in the Earth Galleries Volcanoes and Earthquakes exhibition (Figure 17A).

**Figure 17.**
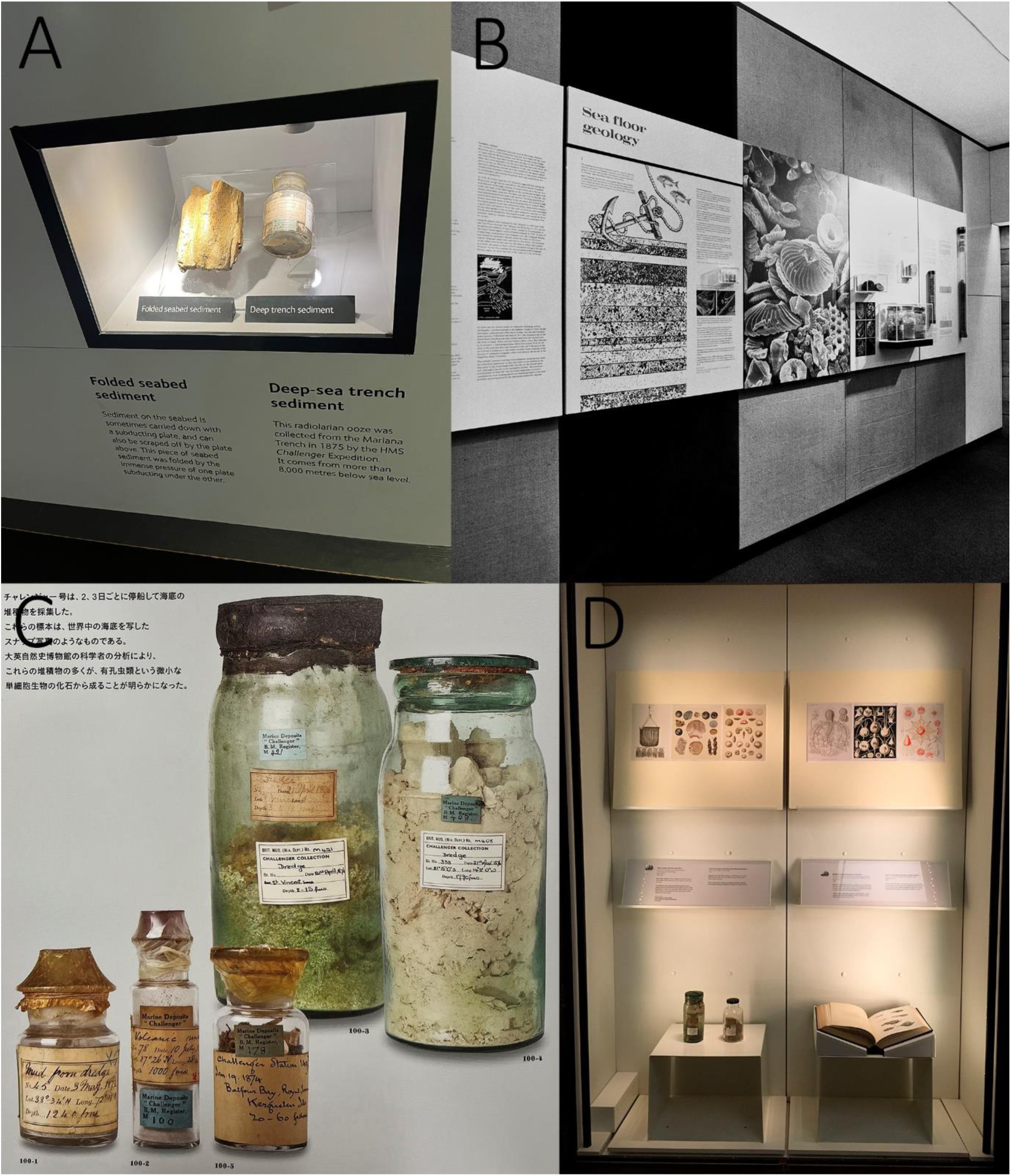
Exhibits. A. Earth Galleries Mariana Trench sample (NHMUK M.289(2)). B. 100^th^ Anniversary temp exhibit showing three glass jars on wall. C. Glass jars from Touring Treasures Exhibit (National Museum of Nature and Science, Tokyo 2017, p. 138). D. 150^th^ Anniversary exhibit in Images of Nature Gallery.

#### Figured Challenger specimens on display boards

It is not possible to locate precise dates for when these specimens were on display or the precise location but the specimens figured in Murray and Renard (1891) are still on their standard British Museum (Natural History) display board. These are corals (Figure 13), pumice, manganese nodules, micrometeorites, sharks teeth (Figure 16), smaller whale parts such as tympanic bones and detailed drawings of slides.

#### 100th Anniversary Celebrations 1973

A note in the collection suggests that some sediment was loaned for an exhibition at Nova Scotia Museum, Canada in 1973. Davis (1973) describes some materials donated to the same museum by the scientists on the Challenger when they stopped off there after their first traverse of the Atlantic in 1874 and presumably these were also included in the exhibition. To mark the 100^th^ Anniversary of the start of the expedition The British Museum (Natural History) held a temporary exhibition ‘The Voyage of HMS Challenger and the Birth of Oceanography’ in the gallery space near to the entrance of the General Library from February 1973 to 10 May 1974. Full details are provided in the Natural History Museum Archives (DF700/2/2, DF/EXH/700/2/2, DF/PH/4/31) but most of the correspondence relates to displayed items borrowed from 12 different institutions. The National Maritime Museum lent a deck plan of Challenger that it would appear from correspondence, was later replaced by a copy. The Science Museum lent most of the equipment including 3 Baillie Sounding rods but no weight, a sextant and two microscopes. The Royal Scottish Museum provided a bust of Wyville Thompson, prints of artwork, charts and a model of The Challenger. There were 6 display cases: Origins, The Ship, Laboratories and instruments, Dredging and Sounding, Personnel and Results. Staff listed as developing the exhibition include J.D.H Wiseman, A.L. Rice, M. Belcher, C.R. Hill as well as K.R.C. Kempe and H.A. Buckley. Items from John Murray’s Collection at the museum included a Miller-Casella thermometer with its case and a hydrometer (M9787). Though not listed in the archives, three rock bottles in a small case can be seen on black and white prints of a panel on Sea Floor Geology (presumably related to the Results display case) along with some manganese nodules (Figure 17b). Murray’s diary and the ship’s log from the Murray Library at the museum were also displayed. A short note in the archive suggests that some basalt and its Perspex case was stolen on 8/3/74 but the numbers given (26-256-10-4) do not conform to any museum numbering system and Murray’s collection does not list any basalt.

#### A world tour

From March 2017-January 2022 The Natural History Museum sent out various Murray Collection items as part of its touring exhibition entitled ‘Treasures of the Natural World’. These are illustrated in the exhibition catalogue for the first exhibition in 2017 at the National Museum of Nature and Science, Tokyo, Japan (18/03/2017 - 11/06/2018) and include various sediment bottles as well as a Miller-Casella thermometer and its brass case (Figure 17c). The exhibition then moved on to The ArtScience Museum, Singapore (22/11/2017 - 29/04/2018) followed by The National Chiang Kai-Shek Memorial Hall, Taipei, Taiwan (28/06/2018 - 19/09/2018) and Musee de la Civilisation, Quebec City, Canada (15/05/2019 - 05/01/2020). A slight hiatus caused by the disruption due to the Pandemic saw it installed at Melbourne Museum, Australia (12/06/2021 - 18/01/2022) before returning to the Natural History Museum later in 2022.

#### 150th Anniversary

A celebratory temporary exhibit was installed in the Images from Nature Gallery from (08/07/2022-12/01/2023) and included some sediment bottles from Sir John Murray’s Collection (Figure 17d).

### Future of the collection

The near future of the collection is a move to the new Natural History Museum storage facility at Reading Thames Valley Science Park at the end of the decade. Preparations will include restorage as well as any conservation work needed on the glass or labelling associated. The collection will also be re-ordered so that similar sizes of glassware and boxes will be kept together to give a more efficient volume of storage. Images of each item will be released gradually on the NHM Data Portal (https://data.nhm.ac.uk/) as the collection is further documented in preparation for the move. Risks of disassociation are greatly reduced as the dataset presented here will allow tracking of individual items. Early versions of this dataset have already facilitated greater access to this collection for current projects on dating of sharks teeth and preparation of funding applications for studying micrometeorites. It is hoped that by listing the contents and detailing the methods of collecting, that further use of the collection will be encouraged. This dataset has wider application in that it can help illuminate other Challenger sediment collections elsewhere, particularly the list of “non-station” deposits documented here providing a wider applicability to studies of Challenger Sediments away from the traditional station deep sea sediment related model. This dataset also allows better association of future works as opposed to the situation where individual digital records covered a wide range of sometimes over 100 different items. Material from this collection is available for loan subject to the Natural History Museum’s loan regulations, for either individual research projects but also for exhibitions. The collection remains topical and has enormous potential for relevant research in climate, carbon sink or biodiversity change projects. One novel use could be the documentation of sedimentary DNA (*sed*aDNA) from relevant samples (Armbrecht et al. 2019) although baseline/background documents such as the current study will be vital in evaluating whether the techniques used on the Challenger are able to provide relevant results. These have real potential for providing hidden details of biodiversity. Climate change studies such as those using Foraminifera have already proven to deliver interesting results and could be expanded to include the plankton samples not covered by this paper. This is particularly relevant with a planned 2025 Promare sponsored cruise of the autonomous vessel Mayflower 400 (https://promare.org/) set to cover the route of the first traverse of the Atlantic by H.M.S. Challenger in 1874. Van Ginneken et al. (2024) have shown the importance of micrometeorite collections for evaluating past environments and changes to our atmosphere over time and the Murray Collection certainly contains many examples that are yet to be released from the sediment. CT scanning and other new techniques such as machine learning can provide excellent non-destructive methods for analysing these sediments (eg Zarkogiannis et al. 2025). Nodules are subject to interesting debates on the ethics of deep sea mining. This collection provides a unique set of samples that will never be able to be collected again but could be used as a baseline for future work. Finally, the sediment collections from the Challenger Expedition provide opportunities for historical studies in aspects such as early exploration techniques, the long term storage of collections in glassware or colonial associations. In all, the detailed dataset and illustrations presented here can lead to richer collaborations, stronger science and ease of repeatability.

## Acknowledgments

Dave Smith (NHM Data Management) helped with Axiell Emu digitisation workflows and volunteers Pelham Miller and Jamie Rolt with data capture relating to the slide collection. Tom Wood and Stergios Zarkogiannis, University of Oxford are thanked for their helpful discussions particularly around the collecting techniques employed on H.M.S. Challenger. Prof Gillen Wood (University of Illinois, USA) kindly gave us a preview of details of his forthcoming book on Challenger. Katie Collins (NHM Fossil Mollusca) gave information about molluscan collections offshore New Zealand. Brian Rosen (NHM Science), Andrea Hart and Antony Stevens (NHM Library and Archives) provided information about the 100^th^ and 150^th^ anniversary exhibitions. Peter Grugeon and Jonathan Jackson, NHM Photo Unit carried out all the photography for the figures. Finally, the Savoy Café, Merton Road kept us fed on Challenger Thursdays.

## Data Availability Statement

The associated dataset is available at https://doi.org/10.5519/50h9e977.

## Appendices

### 1.1 List of samples labelled as having been rebottled

M.18(5) 13^th^ February 1956 M.21(10) August 1958

M.414(2) retubed March 1959.

M.114(3), Transferred from cracked bottle, April 1959

M.373(22). Rebottled May 1959.

M.339(7) Label inside tube says ‘retubed Oct. 1959

M.339(8) reboxed October 1959

M.339(9) reboxed October 1959

M.339(11) retubed October 1959

M.44(1), November 1959

M.353(49) Retubed 1959.

M.357(37,38) Retubed 1959. Contaminated 1966.

M.367(8) Rebottled Jan 1960. Section prepd.

M.25(15) rebottled 29.8.1974. M.372(1) Rebottled May 1975.

M.372(19). rebottled May 1975.

M.339(10), original butter jar found broken 1.10.1993.

### 1.2 Samples are absent from these 99 Stations listed in Appendix II of Tizard et al. (1895). Details in the *Deep Sea Deposits* volume (Murray and Renard, 1891) suggest that samples were not retained for various reasons from the stations listed in bold

**I, I(A,B,C,D)**, **II(C,K**), **III, IV**, **VA**, **VII**, **VII(G,H,J,K**,**N**,**Q**,**S**,**U,V**), **4**, **6**, 23A, **32(C,E,F,G)**, **33(A,B), 34**, **36**, **41**, **43**, **48**, **49**, **52A**, **55A**, **56A, 57**, **77**, **84**, **93(D,E)**, **96**, **99**, **100**, **103**, **105**, **109(A,B)**, **113**, **113(B,C)**, **114**, 122(B,C), **135(A,B,D,E,G)**, **136**, 145, 145A, **148, 148A**, 161, 163(A,B), **164(C,D,E)**, 166(**A**,C), 167A, **170**, **170A, 171**, **173**, **174A**, 185A, **197**, **200**, **201**, 203, **208**, **212**, 233C, 236A, 301, **312**, **314A**, 315, 316, **340**, **349, 350, 351, 352**.

### 1.3 Column descriptions for dataset

First Tab - Items

*Sort number*. This is a sequential number added to aid sorting the dataset by chronological date of collection.

*irn.* This internal record number serves to aid retrieval of records for the NHM Collections Management System which is currently in Axiell Emu but will be migrated to a new bespoke system sometime in 2025. This number also appears on the NHM Data portal as well as the BGIF portal, to which these records are also delivered.

*NHM registration number.* The majority of these are John Murray (M) numbers. Originally only one number was assigned to a set of items mostly from the same collecting site and collected on the same day. An audit of the entire collection has allocated subnumbers that appear here in brackets to help delimit individual items covered by these original “M” numbers. Other prefixes include a number based system used for the petrology collections (eg BM.1920, 669(19))

*M base number.* This unformatted M number column has been included to aid sorting by M number.

*Station number*. These have been included when possible, although some sampling happened at sites where Station Numbers were not allocated, particularly for shallower deposits near to shore or in harbours. When the associated dates suggest that sampling occurred between numbered stations a range has been given.

*Verbatim label*. This is a transcription of all label data on items. Eventually the intention is to provide an image of each item on the NHM data portal so the labels will be visible as images as well as transcriptions.

*Sounding number*. Appendix II of the Narrative to the cruise (Tizard et al. 1885) gives a sounding number associated with each collecting site. When a station number has not been allocated, this field is left blank. As noted in the main text, the sounding numbers in this appendix and the sounding numbers recorded verbatim on the labels do not correspond as the verbatim label numbers are derived from Murray (1876).

*Method of collecting*. This is one of or a variant of: Sounding, Dredge, Trawl, Surface Net, Tow Net, Beach Sand, Anchor Mud.

*Type of container.* This has been standardised to start with the composition and type and of container followed by approximate size and a comment on the type of lid if applicable eg. Cardboard box, small, glass lid.

*Date collected.* This has been included on both of the first two tabs as sometimes more than one date has been included for items grouped under one M number. There are also variations on dates allocated between original bottles and preparations eg slides.

*Depth in fathoms*.

*Notes*.

Second Tab – List of sites

*Sort number*. Sequential number used to aid sorting in chronological order.

*IRN Catalogue of first (base) record*. Internal record number for first specimen record attached to collecting site.

*Base registration number.* Alphanumeric NHM specimen registration number.

*Original record sample quantity details*. Legacy data from when each M number was registered at the NHM to represent multiple specimens. Usually refers to numbers of bottles, tubes, slide or boxes.

*Nature of Bottom*. Details of type of sediment verbatim from Tizard et al. (1881) Appendix II.

*Date Visited*

*Station number*

*Water body*. Major ocean bodies including standard internationally recognised subdivisions.

*Extra land mass based detail.* For sites near to land eg from harbours or offshore islands.

*Lat – given*. As per Tizard et al. (1881) Appendix II.

*Long – given*. As per Tizard et al. (1881) Appendix II.

*Georef Lat.* Georeferenced latitude based on maps present in Tizard et al. (1885), Murray and Renard (1891) and Murray (1885).

*Georef Long.* Georeferenced longitude based on maps present in Tizard et al. (1885), Murray and Renard (1891) and Murray (1895).

*Site Summary Data.* Internal NHM collections management system summary string representing Site details.

*NHM Site irn.* Internal NHM collections management system internal record number representing Site.

### 1.4 List of samples affected by 6/3/1966 flood

M.194(12-20, 23-26, 31, 35, 54-63), stn 160;

M.306(23), stn 241,

M.320(1-11, 15, 17-20) stn 256,

M.353(51, 70, 77), stn 281,

M.355(2), stn 283,

M.357(4, 28, 38, 83), stn 285,

M.382(4) stn 309,

M.384(5), stn 310,

M.385(2, 3), stn 311, M.383(1), stn 309a,

M.384(5,) stn 310,

M.385(2,3), stn 311,

M.386(1), stn 313,

M.387, stn 314,

M.406(7), stn 334,

M.407(3), stn 335,

M.420(1), St Vincent (no station number)

